# Nanoparticle Albumin-Bound Paclitaxel Targets Pulmonary Neutrophils

**DOI:** 10.64898/2026.09.27.754839

**Authors:** Aparajeeta Majumder, Soomin Jeong, Marco E. Zamora, Nicolas Marzolini, Yufei Wang, Jia Nong, Taylor Brysgel, Priyal Patel, Michael H. Zaleski, Jichuan Wu, Collin T. Stabler, Liam S. Chase, Zhicheng Wang, Ed Morrisey, Vladimir R. Muzykantov, Laura Ferguson, Jacob S. Brenner, Oscar A. Marcos-Contreras, Jacob W. Myerson

**Author notes:** Corresponding authors Jacob S. Brenner, Oscar A. Marcos-Contreras, Jacob W. Myerson.

## Abstract

Nanoparticle albumin-bound paclitaxel (Abraxane) was among the first clinically approved nanomedicines and is a first-line chemotherapeutic. It enhances therapeutic efficacy and reduces side effects of paclitaxel. Despite its extensive clinical use, mechanisms of Abraxane’s therapeutic effects are the subject of ongoing study. This work posits a new mechanism for Abraxane’s effects in lung cancers and metastasis. Tracing the albumin component of Abraxane with radioisotopes or fluorophores, we find that Abraxane homes to the lungs. Abraxane’s lung tropism is elevated by inflammation and is eliminated in complement protein C3 knockout mice, showing that complement pathway drives the uptake. Flow cytometry and histology show that neutrophils are the dominant cell type taking up Abraxane in the lungs. We demonstrate Abraxane’s tropism to pulmonary neutrophils in mice and in human donor lungs. We note previously undocumented side effects associated with Abraxane’s lung tropism, including pro-thrombotic and pro-inflammatory responses in acute lung inflammation. We show that Abraxane’s efficacy against lung metastases is affected when lung uptake is eliminated by C3 knockout. Our results provide insight into why Abraxane is effective in lung cancer and metastases. These findings may be used to improve outcomes in patients receiving Abraxane and help design next-generation cancer nanomedicines.

**Teaser:** Nanoparticle albumin-bound paclitaxel, widely used for treatment of non-small cell lung cancer and metastases arising from breast and pancreatic cancer, was found to target neutrophils in the lungs.

## Introduction

Nanoparticle albumin-bound paclitaxel (Abraxane) is among the most clinically impactful nanomedicines.(*1–3*) It is a first-line medicine for multiple cancers. It was initially approved for metastatic breast cancer in 2005(*4*) and has subsequently reached the clinic for non-small cell lung cancer,(*2, 5*) metastatic pancreatic cancer,(*6*) and PD-L1-positive metastatic triple-negative breast cancer.(*7, 8*) This study proposes a new mechanism controlling Abraxane’s biodistribution, with implications for its chemotherapeutic effects in the lungs, an important niche for all of these cancers.

Abraxane represents a success story for cancer nanomedicine traversing the history of the field. It was first developed over three decades ago, employing albumin as a carrier complexed with paclitaxel as a drug component, with hydrophobic interactions between the two components forming ∼130 nm diameter nanoparticles (Fig. 1a).(*2, 9–11*) Abraxane’s structure complexes paclitaxel with a biocompatible protein, altering its pharmacokinetics and limiting its toxicity.(*12, 13*)

**Figure 1.**
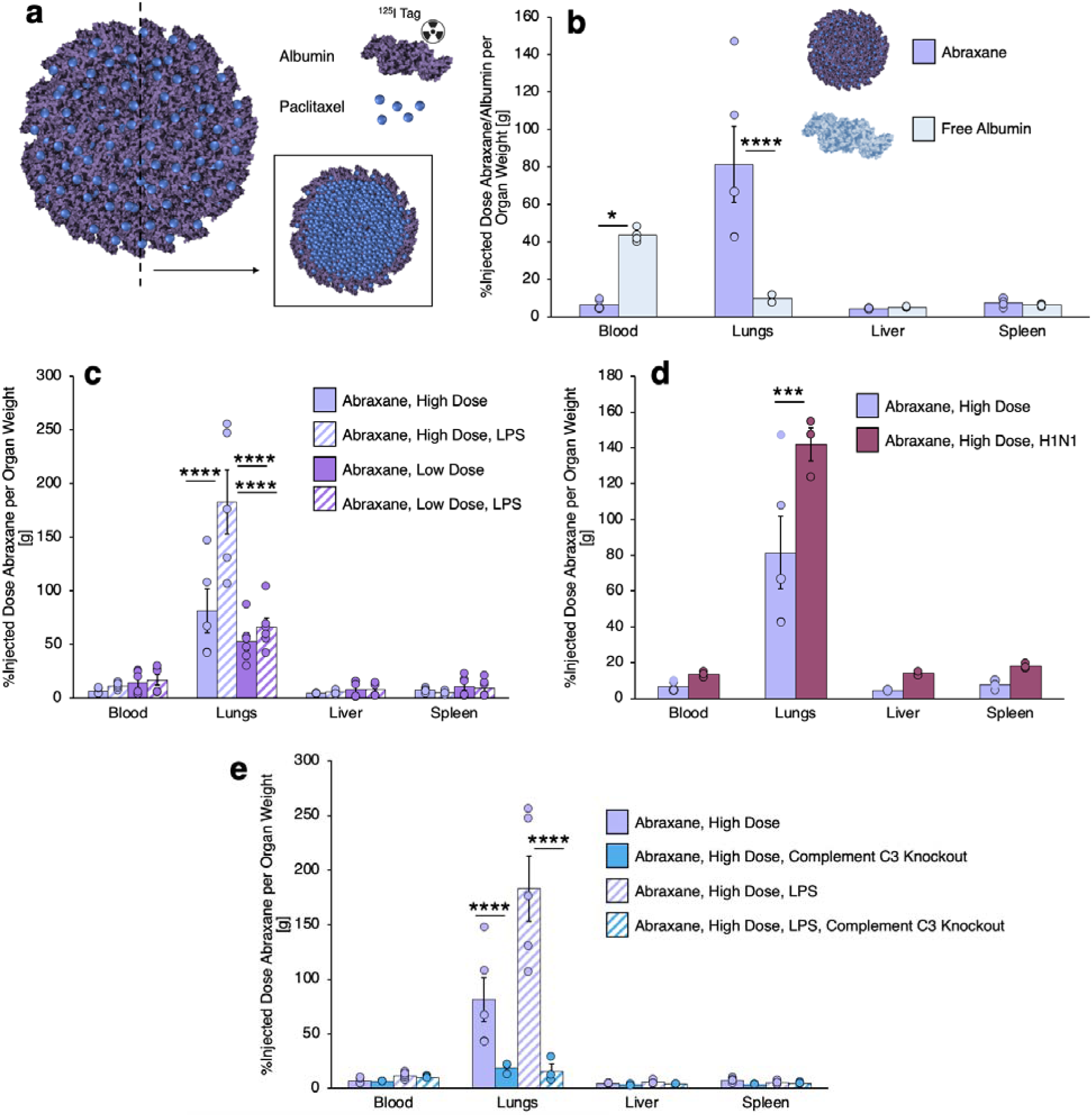
**Biodistributions of radiolabeled albumin in Abraxane**. (a) Schematic of Abraxane, indicating radiolabeling of the albumin component for biodistribution experiments. Inset: cross-section showing the core-shell structure of Abraxane. (b) Biodistributions of 5 mg/kg Abraxane (n=4) and albumin alone (n=4) in naïve mice, showing high lung uptake of Abraxane. (c) Biodistributions of Abraxane in naïve mice (5 mg/kg, n=4 and 1 mg/kg, n=6) and in mice treated with intravenous LPS (5 mg/kg, n=5 and 1 mg/kg, n=6), showing increase in high-dose Abraxane lung uptake induced by acute inflammation. 5 mg/kg Abraxane data in naïve mice is reproduced from (b) for comparison to the other data sets. (d) Biodistributions of 5 mg/kg Abraxane in mice afflicted with H1N1 influenza (n=3), showing increased lung uptake in influenza-affected mice. 5 mg/kg Abraxane data in naïve mice is reproduced from (b) for comparison. (e) Biodistributions of 5 mg/kg Abraxane in naïve C3 knockout mice (n=3) and C3 knockout mice treated with intravenous LPS (n=3), showing that lung uptake of Abraxane depends on complement. Biodistributions of Abraxane in wild-type mice under corresponding conditions are reproduced from (c) for comparison. *p=0.01, ***p=0.0009, ****p<0.00001 using two-way ANOVA with Sidak’s correction for multiple comparisons.

Despite the lasting success of Abraxane, mechanisms by which it improves paclitaxel’s therapeutic effects are a topic of ongoing study. Tracing eluted paclitaxel does not confirm the hypothesis that Abraxane permits enhanced tumor uptake or prolonged blood retention relative to prior Taxol formulations.(*14–16*) More recent work has traced the albumin component of Abraxane, considering how albumin impacts uptake of Abraxane in tumors, including via endogenous albumin receptors and albumin-binding proteins.(*17*) But the literature doesn’t yet show a clear organ- or cell-specific targeting mechanism for Abraxane.

The biodistributions of albumin nanoparticles can be sensitive to their structure.(*18, 19*) The way albumin is arranged in or on nanoparticles can predict accumulation in the lungs, especially in settings of acute lung inflammation. Generally, dense packing of proteins like albumin in nanoparticles, including via hydrophobic interactions, predicts nanoparticle uptake in leukocytes in the pulmonary vasculature.(*20*) Controlling the physicochemical properties of nanoparticles, such as the structural arrangement of albumin in Abraxane, is a way to formulate nanoparticles with tropism for specific organs and cell types without the need for affinity moieties.(*21–23*) That is, the structure of a nanoparticle can give it targeting behaviors, even if the components of the nanoparticle don’t have targeting behaviors themselves.

Here, we reevaluate Abraxane through the lens of structure-based physicochemical tropism. We hypothesized that similarities between the characterized structure of Abraxane and the structures of recently developed albumin nanoparticles with physicochemical lung tropism would reveal previously undetected lung tropism for Abraxane. Evaluating Abraxane’s biodistribution using albumin radiolabeling, we find that the albumin component of Abraxane concentrates in the lungs after intravenous injection, with acute inflammatory conditions increasing lung uptake to levels >20% of injected dose. We demonstrate that this lung uptake is dependent on the complement pathway immune response to Abraxane, with lung tropism being completely ablated in complement C3 knockout mice. We show that Abraxane’s lung uptake is achieved by high selectivity for pulmonary neutrophils. Abraxane’s lung tropism yields previously undocumented inflammation-dependent side effects, including thrombosis and increased IL-6 concentration in lungs affected by prior acute inflammation. Abraxane’s lung tropism and selectivity for pulmonary neutrophils are reproduced in *ex vivo* studies with perfused and inflated human lungs. Abraxane’s efficacy against lung metastases is reduced by ablation of its lung uptake in C3 knockout mice. Our work therefore presents a new perspective on Abraxane: It is a lung targeting agent whose efficacy is linked to delivery to pulmonary neutrophils, a cell type with important roles in metastasis that has drawn interest as a tool for cancer targeting(*24–27*).

## Results

### Abraxane targets the lungs

Noting literature demonstrating albumin nanoparticles with lung tropism(*18, 19*), especially in acute inflammatory disorders, this portion of the results presents experiments tracing Abraxane’s biodistribution in naive mice and mice with different forms of acute lung inflammation.

Studies of Abraxane’s biodistribution have typically traced the therapeutic component of the nanoparticle, paclitaxel, with mass spectrometry techniques(*14, 28*). Recent work has discovered new behaviors or Abraxane while tracing the albumin component of the drug(*17*). Our studies take this latter tact, labeling albumin in Abraxane with ^125^I radiotracer or AlexaFluor 647 fluorophore (Figure 1a). Applying either label to Abraxane from pharmacy stock, we find nanoparticle diameters of ∼120-130 nm, with labeling inducing no significant changes in size, concentration, or polydispersity of Abraxane, as measured by nanoparticle tracking analysis and dynamic light scattering (Supplementary Figure 1).

Radiolabeled Abraxane was traced in mice after injection as a bolus at 5 mg/kg drug dose. As a control, we traced radiolabeled (^125^I) human albumin at the same bolus dose. Our results confirm that the albumin in Abraxane nanoparticles does not distribute in the body like albumin alone. In biodistributions 30 minutes after injection, Abraxane uptake in the lungs is significantly elevated relative to albumin alone (Figure 1b). The isolated albumin concentration in the lungs is <2% of injected dose. As expected, isolated albumin maintains a high concentration in the blood. But Abraxane achieves a lung concentration of ∼80% of dose per gram tissue (∼10% of total dose retained in the lungs). Concurrently, the blood pool concentration of Abraxane is significantly diminished relative to albumin alone. Among other tested organs, there were no significant differences between Abraxane retention and albumin retention (Supplementary Figure 2a-b).

Since prior studies have demonstrated albumin nanoparticles with unique tropism for inflamed lungs vs. naive lungs, we traced Abraxane’s biodistribution in mice with bacterial lipopolysaccharides (LPS)-induced inflammation. At 5 mg/kg bolus dose, Abraxane uptake in the inflamed lungs is significantly elevated relative to that in naive lungs (Figure 1c, Supplementary Figure 2a). In LPS-inflamed lungs, Abraxane achieves concentrations of ∼180% of dose per gram lung tissue (∼20-25% of total dose retained in the lungs). Abraxane’s selectivity for inflamed lungs is dose-dependent. At a 1 mg/kg bolus drug dose, Abraxane concentrates in the lungs at ∼50-70% of dose per gram lung tissue, with no significant differences between naive and inflamed lungs. In naive lungs, uptake of 1 mg/kg Abraxane, as a fraction of injected dose, does not differ significantly from uptake of 5 mg/kg Abraxane.

To study high-dose Abraxane selectivity for inflamed lungs in a more clinically relevant mouse model, mice were infected with H1N1 influenza(*29*). After inhalation of H1N1, Abraxane uptake in the lungs significantly increases (Figure 1d, Supplementary Figure 2c). In influenza-affected lungs, 5 mg/kg Abraxane retains at a concentration of ∼140% dose per gram tissue (∼15-20% of dose). Influenza does not change Abraxane retention in other tissues.

Our previous work showed that protein nanoparticle tropism for the lungs can be dependent on opsonization of the nanoparticle by complement pathway proteins(*18, 20, 30, 31*). Specifically, we found that nanoparticles formed by hydrophobic interactions between proteins accumulate in pulmonary neutrophils in a manner that can be blocked by complement inhibition. Hypothesizing that Abraxane, a nanoparticle formed by hydrophobic interactions between albumin and paclitaxel, would behave similarly, we traced Abraxane in complement protein C3 knockout mice. In both naive C3 knockout mice and C3 knockout mice treated with intravenous LPS, Abraxane uptake in the lungs was eliminated (Figure 1e, Supplementary Figure 2d).

### Abraxane targets pulmonary neutrophils

Recent work from our group and others has demonstrated a dominant role for neutrophils in a variety of strategies for nanoparticle delivery to the lungs, particularly in acute inflammatory conditions(*18, 20, 30, 31*). For many lung-targeting nanoparticles, we have established a connection between complement activation and neutrophil tropism. We hypothesized that Abraxane’s complement-dependent tropism for the lungs reflects a tropism for the abundant neutrophils in the pulmonary vasculature. This section of the results presents histological and flow cytometry evidence for Abraxane’s selectivity for neutrophils in the lungs.

Alexa Fluor 647-labeled Abraxane was given as a 5 mg/kg bolus to naive mice and mice treated with intravenous LPS. Sections of the lungs were stained with DAPI, FITC-labeled CD45 antibody to identify leukocytes, and Alexa Fluor 594-labeled PECAM antibody to identify endothelial cells. Confocal imaging of lungs affected by LPS shows abundant Abraxane signal, primarily in puncta (Figure 2a, Supplementary Figure 3, Supplementary Movie 1). Focusing specifically on Abraxane colocalization with CD45, we observe evidence of Abraxane homing to leukocytes (Figure 2b, Supplementary Figures 4-5, Supplementary Movie 2). We obtain similar histology results in LPS-affected and naive lungs (Supplementary Figure 6).

**Figure 2.**
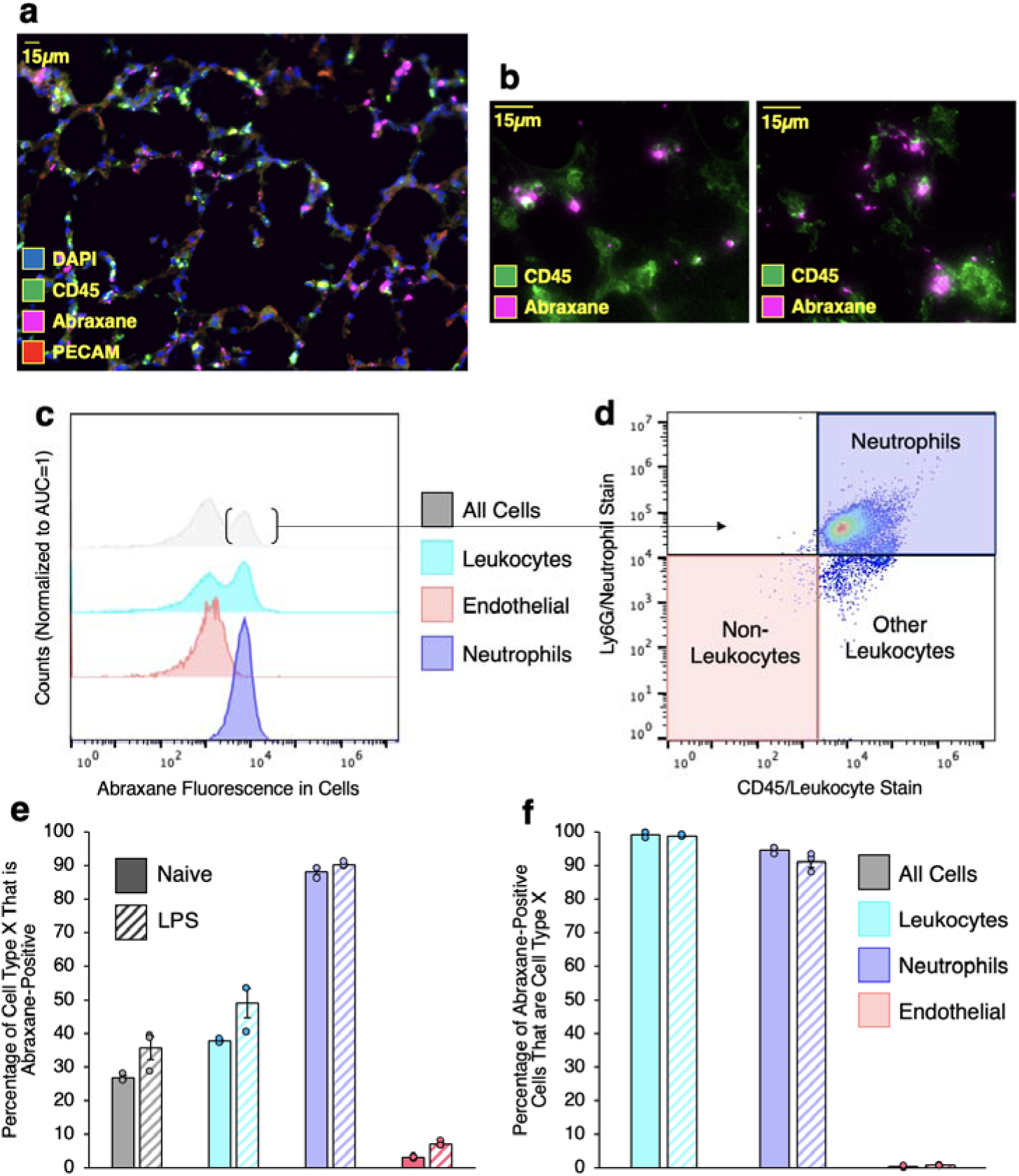
**Histology and flow cytometry assessment of Abraxane distribution to different cell types in the lungs**. (a) Lung micrograph with staining for DAPI (blue), CD45/leukocytes (green), and PECAM (red). Lungs are from mice treated with intravenous LPS prior to dosing with Alexa-647 labeled Abraxane (pink) (n=2 mice). (b) Higher magnification of lung micrographs with identical conditions to (a), isolating Abraxane and CD45 signals to show colocalization. (c-d) Flow cytometry data characterizing cell suspensions prepared from lungs of mice treated with intravenous LPS prior to dosing with Alexa-647 labeled Abraxane. Parentheses indicate the fraction of all recovered cells that were categorized as Abraxane-positive. Data in (d) show CD45 and Ly6G staining among Abraxane-positive cells. (e-f) Summary of flow cytometry data showing that; (e) ∼90% of lung neutrophils are Abraxane-positive and; (f) >95% of Abraxane lung uptake is attributable to leukocytes and >90% of Abraxane lung uptake is attributable to neutrophils in both naïve and intravenous LPS-treated mice (n=3 mice for each condition).

To quantitatively assess Abraxane uptake in pulmonary leukocytes, single cell suspensions were prepared from mouse lungs after dosing with a 5 mg/kg Abraxane bolus. The cells were stained to identify endothelial cells, total leukocytes, and neutrophils (Supplementary Figure 7). Abraxane fluorescence was quantified in each cell type, as well as the total population of cells recovered from the lungs (Figure 2c). A subpopulation of all recovered cells, and of the recovered leukocytes, has elevated Abraxane fluorescence. Isolating endothelial cells, we find no Abraxane-high cells. But when we isolate neutrophils, we find nearly all cells are Abraxane-positive. Similar results are obtained for LPS-challenged and naive lungs (Supplementary Figure 8). Likewise, focusing on the total population of Abraxane-positive cells, we find that the vast majority are CD45/Ly6G double positive, indicating neutrophils (Figure 2d).

Summarizing the flow cytometry data; 1) >90% of pulmonary neutrophils are Abraxane-positive, compared to <5% of endothelial cells (Figure 2e); 2) >90% of Abraxane-positive cells are neutrophils, and >95% are leukocytes (Figure 2f). To show that Abraxane’s selectivity for leukocytes in general and neutrophils in particular reflects uptake in individual cells, we re-analyzed our flow cytometry data to include cell aggregates. We find no difference in Abraxane selectivity for neutrophils when comparing data including cell aggregates with data isolating single cells (Supplementary Figure 9).

### Abraxane has thrombotic and inflammatory side effects in acute lung inflammation

In mice with LPS-induced acute lung inflammation, we noted intermittent gross side effects of 5 mg/kg Abraxane, including sluggish movement and mortality (two of 22 mice receiving 5 mg/kg Abraxane following intravenous LPS died within 30 minutes of Abraxane dosing). Abraxane’s lung targeting is dependent on complement and sensitive to acute lung inflammation, so we hypothesized that Abraxane may exert additive proinflammatory effects in the lung vasculature in settings of pre-existing acute inflammation.

Lung sections were prepared from mice treated with Abraxane boluses. Masson trichrome staining showed differences in the pulmonary vasculature of mice receiving Abraxane after intravenous LPS, compared to LPS-affected mice not receiving Abraxane and to naive mice, with or without Abraxane treatment (Figure 3a, Supplementary Figures 10-11). Particularly in larger vessels, occlusive thrombi were present in mice treated with Abraxane after intravenous LPS. Blinded scoring of Masson trichrome staining shows that risk of pulmonary thrombosis is significantly elevated in mice treated with Abraxane following acute lung inflammation (Figure 3b, Supplementary Figure 12).

**Figure 3.**
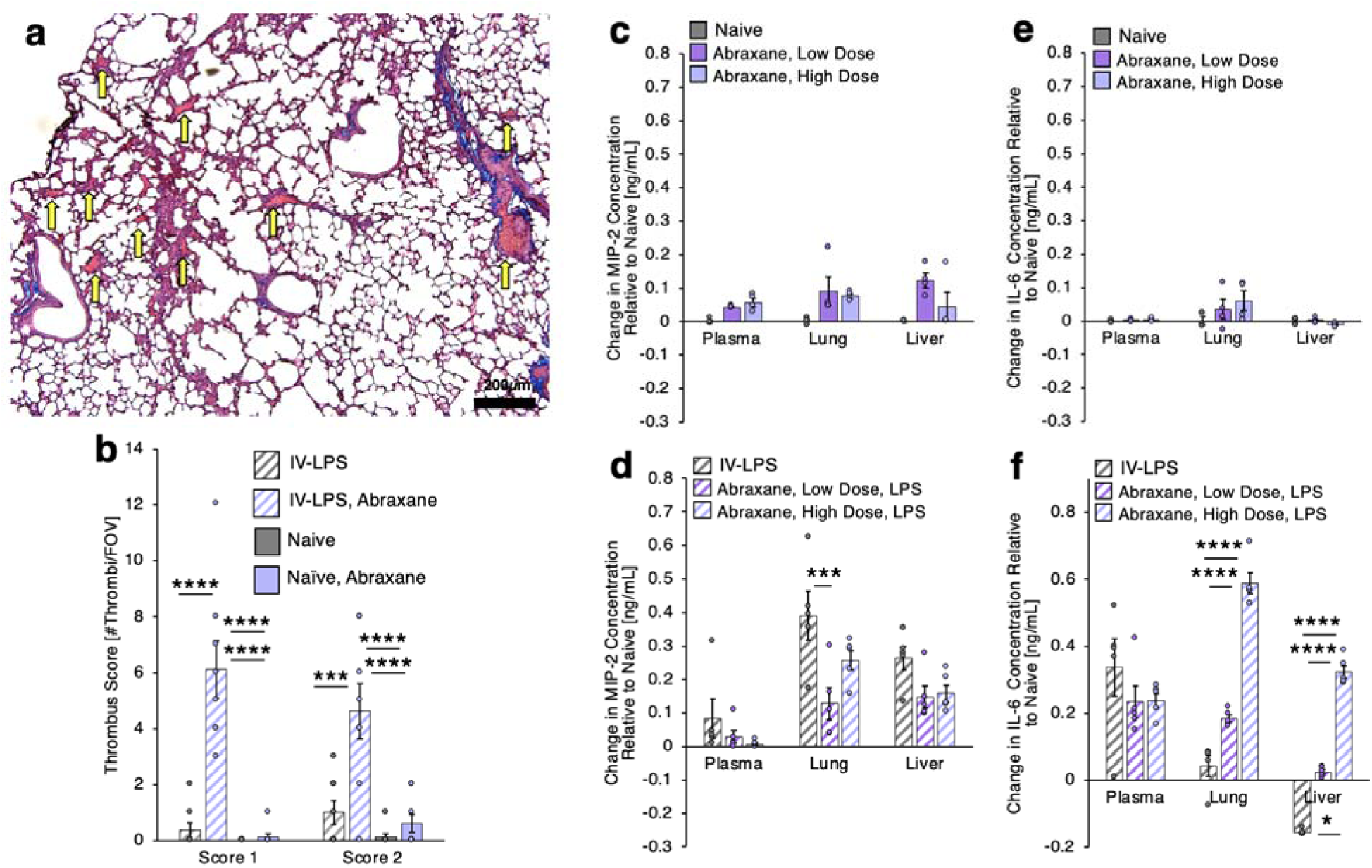
**Side effects of Abraxane lung uptake**. (a) Masson trichrome-stained lungs of mice treated with intravenous LPS prior to dosing with Abraxane (5 mg/kg), showing thrombi (yellow arrows). (b) Masson trichrome lung histology data as adjudicated by two blinded scorers, showing elevated lung thrombosis in mice treated with Abraxane following intravenous LPS (n=8 micrographs over 3 mice per condition, ****p<0.00001, ***p=0.0002). (c-d) Macrophage inflammatory protein 2 (MIP-2) levels, with naïve averages subtracted, in mice treated with 1 mg/kg or 5 mg/kg Abraxane, with or without prior intravenous LPS. Data show elevated MIP-2 in the lungs and liver after intravenous LPS, with 1 mg/kg Abraxane treatment blunting the lung elevation. (e-f) Interleukin 6 (IL-6) levels, with naïve averages subtracted, in mice treated with 1 mg/kg or 5 mg/kg Abraxane, with or without prior intravenous LPS. Data show elevated IL-6 in plasma and depressed IL-6 in the liver after intravenous LPS. In a dose-dependent manner, IL-6 in the lungs and livers of LPS-treated mice is elevated by Abraxane (in c-f, n=4 for all naïve conditions and n=5 for all LPS-treated conditions, *p=0.01, ***p=0.0006, ****p<0.00001). All statistics obtained using two-way ANOVA with Sidak’s correction for multiple comparisons.

Our recent work has characterized modulation of inflammatory signaling as part of immune interactions with nanoparticles in settings of preexisting acute inflammation(*18, 32, 33*). We have noted sensitivity of macrophage inflammatory protein 2 (MIP-2) and interleukin 6 (IL-6) to nanoparticles with complement-mediated neutrophil tropism and to solid lipid nanoparticles targeted to the lungs. We therefore quantified plasma, lung, and liver MIP-2 and IL-6 responses to Abraxane (Figure 3e-h). MIP-2 is a chemokine associated with neutrophil chemotaxis. It is elevated in the lungs and liver by inflammatory insults(*32, 34*). Abraxane induces a trend towards lower MIP-2 levels in LPS-affected lungs and livers, with a significant effect in the lungs for the 1 mg/kg dose. IL-6 has broad pro-inflammatory effects and is elevated in plasma and in the lungs in a variety of inflammatory conditions(*33, 35*). IL-6 levels in the lungs and livers are significantly altered by Abraxane after intravenous LPS insult. Abraxane induces dose-dependent increases in IL-6 in the lungs and livers of mice with LPS-induced inflammation.

Our assessment of the acute inflammatory and thrombotic effects of Abraxane in preexisting inflammation is not comprehensive, but we note no additional effects in other broad metrics for side effects. Complete blood count analysis shows Abraxane exerting no acute effects on circulating leukocytes, platelets, or red blood cells, in either naive mice or mice treated with LPS (Supplementary Figures 13-15). Ektacytometry confirms the lack of effects on red blood cells, showing erythrocytes having normal mechanical properties after Abraxane treatment (Supplementary Figure 16).

We also probed for effects of 5 mg/kg Abraxane on the outcome of LPS-induced acute respiratory distress in mice. Lung injury was induced in mice with nebulized LPS, resulting in weight loss and development of pulmonary edema, as measured by accumulation of protein and leukocytes in bronchoalveolar lavage fluid (Supplementary Figure 17). We previously found that liposomes with neutrophil tropism can modulate the behavior of neutrophils in acute lung injury, reducing the quantities of protein and leukocytes in bronchoalveolar lavage fluid(*18*). Abraxane exerts no such effects, either on weight loss or on pulmonary edema.

### Abraxane targeting to pulmonary neutrophils is recapitulated in human lungs

The studies described above provide data indicating that Abraxane achieves focal targeting to and effects in the lungs that were not previously documented in extensive preclinical and clinical testing. Acknowledging the potential clinical relevance of this work, the below portion of the results translates our findings in mice to experiments in human lungs.

Lobes from human donor lungs rejected for transplant were prepared for intra-arterial administration of radiolabeled or fluorescent Abraxane (Supplementary Figure 18). After inflation of the selected lobe (Figure 4a, left panel), an artery was cannulated and the lobe was injected with tissue dye. The tissue dye identified; 1) areas of the lobe perfused by the cannulated artery; 2) veins through which material injected into the cannulated artery exited the lobe (Figure 4a, right panel). In radiotracer experiments, ^125^I labeled Abraxane was injected concurrently with ^131^I labeled human albumin. ∼37% of injected Abraxane retained in portions of the lung marked by tissue dye, compared to ∼10% of injected albumin (Figure 4b).

**Figure 4.**
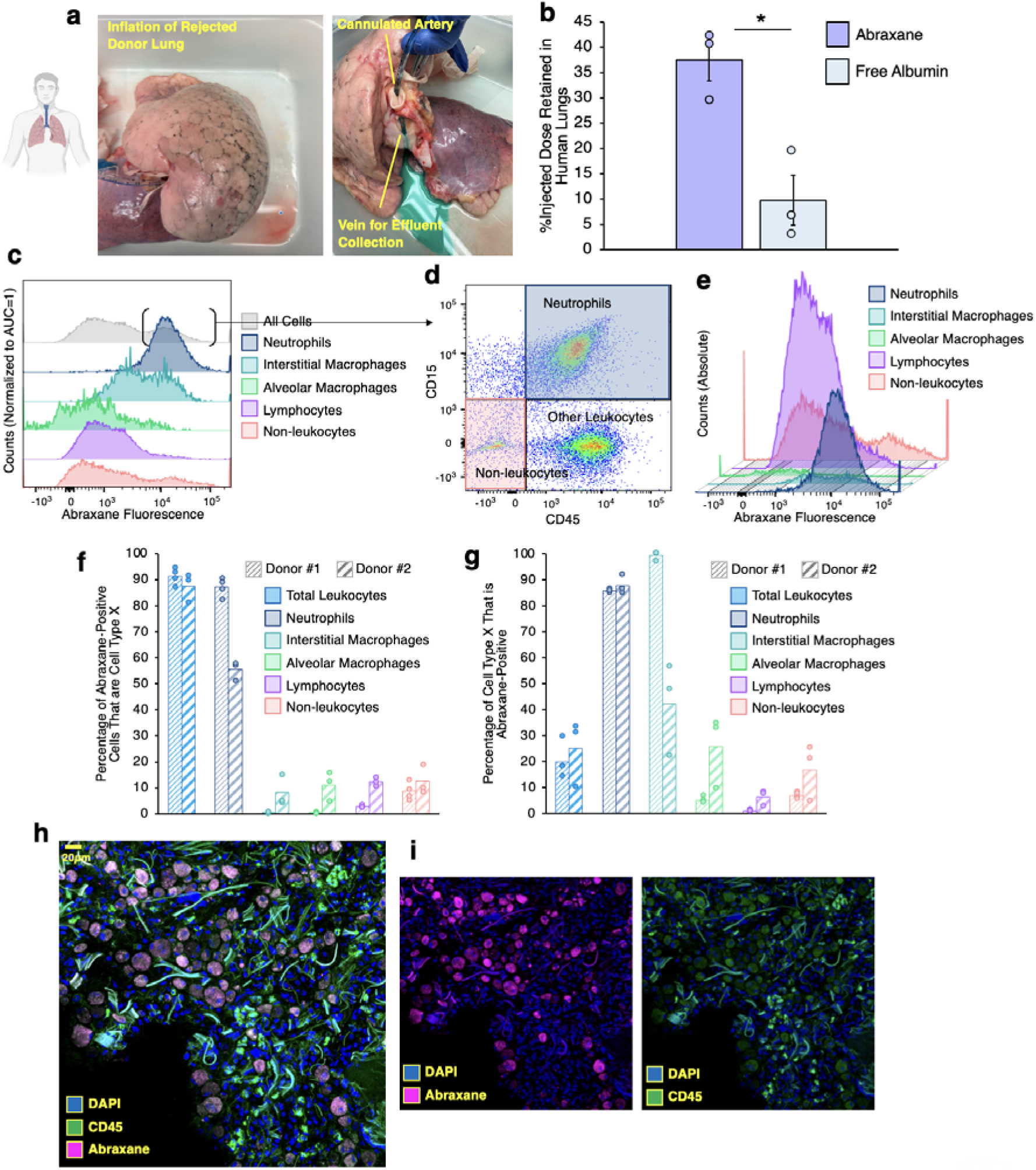
**Neutrophil-driven Abraxane uptake in *ex vivo* human lungs**. (a) Photographs depicting inflation and cannulation of rejected human donor lung lobes. Right: Cannulated arteries were injected with green tissue dye prior to dosing with Abraxane or albumin to mark tissue perfused by the cannulated artery and to identify veins connected to the cannulated artery, from which unbound Abraxane and albumin were collected. (b) Retention of radiolabeled Abraxane vs. radiolabeled albumin in rejected human donor lungs (n=3 lungs). Abraxane was labeled with ^125^I and albumin was labeled with ^131^I, enabling simultaneous tracing of the two in each tested lung. (c-e) Flow cytometry data identifying cell types taking up Alexa-647 labeled Abraxane in human lungs. (c) Abraxane fluorescence intensity in each tested cell type. Parentheses indicate the portion of all cells identified as Abraxane-positive. (d) CD45 and CD15 (identifying neutrophils) staining among Abraxane-positive cells. (e) Data as in (c), but showing the abundance of each tested cell type, indicating high absolute numbers of Abraxane-positive neutrophils vs. Abraxane-positive cells of other types. (f-g) Summary of flow cytometry data from human lungs, indicating data from two separate donors (n=4 tissue samples from donor 1 and n=3 tissue samples from donor 2). (f) Percentage of Abraxane-positive cells that were leukocytes, neutrophils, interstitial macrophages, alveolar macrophages, lymphocytes, or non-leukocytes (n=4 samples from donor 1, n=3 samples from donor 2). (g) Percentage of leukocytes, neutrophils, interstitial macrophages, alveolar macrophages, lymphocytes, or non-leukocytes that were Abraxane-positive. (h-i) Histology data showing Abraxane uptake in leukocytes in human lungs. *p=0.01 by paired t-test.

In experiments with fluorescent Abraxane in human lungs, perfused portions of the lungs were reserved for either flow cytometry or histology. For flow cytometry studies, single cell suspensions were prepared from Abraxane-perfused lung segments, as in mouse studies. To assess Abraxane interactions with leukocytes in the lungs, cell suspensions were stained to identify total leukocytes, neutrophils, interstitial macrophages, alveolar macrophages, and total lymphocytes (see Supplementary Figure 19 for gating strategy).

Quantifying Abraxane fluorescence in each cell type, we find an Abraxane-high population among interstitial macrophages, but, as in mice, nearly all neutrophils recovered from human lungs have high levels of Abraxane fluorescence (Figure 4c). Evaluating stains for leukocytes and neutrophils among all Abraxane-positive cells recovered from the lungs, we find that a majority are neutrophils and nearly all are leukocytes (Figure 4d). Indeed, while both neutrophils and interstitial macrophages have Abraxane-high populations, representing Abraxane fluorescence histograms in terms of the absolute abundance of each cell type shows that the number of Abraxane-high neutrophils dwarfs that of other cell types (Figure 4e). The predominance of leukocytes among Abraxane-high cells is evident in cells from two different donors, despite donor-to-donor variation in the abundance of different cell types (Supplementary Figure 20).

Overall, in lungs from two donors, ∼90% of Abraxane-positive cells were leukocytes. Neutrophils accounted for ∼85% and ∼55% of Abraxane-positive cells in the two tested donor lungs. Differing abundance of other leukocytes competing for Abraxane accounted for the donor-to-donor differences in neutrophil uptake (Figure 4f). When isolating neutrophils among recovered cells, 85-90% of neutrophils were Abraxane-positive in both donor lungs, indicating that Abraxane has uniform neutrophil affinity across donors (Figure 4g). Analysis of flow cytometry data to include cell aggregates indicates neutrophil aggregates may contribute to a small portion of Abraxane uptake in human lungs (Supplementary Figures 21-22). Assessment of endothelial cells and epithelial cells shows low Abraxane uptake. There was donor-to-donor variation in Abraxane uptake in endothelial cells, with ∼40% being Abraxane positive in one donor and <10% in another (Supplementary Figure 23).

Histological analysis confirms the tropism of Abraxane for neutrophils in human lungs. DAPI and CD45 staining show diffuse Abraxane signal in leukocytes with the morphological characteristics of neutrophils and macrophages (Figure 4h-i, Supplementary Figure 24, Supplementary Movies 3-6). Abraxane signal consistently colocalizes with CD45 staining in our data. Co-staining CD45 and CD15 shows that the majority of Abraxane-positive leukocytes in our human lungs histology data are neutrophils (Supplementary Figure 25, Supplementary Movies 7-10).

### Therapeutic effects of Abraxane are affected by lung targeting

Abraxane is a first-line therapy for non-small cell lung cancer. It is also used for metastatic breast and pancreatic cancers. Abraxane’s utility in lung cancer and metastases(*2–8*) led us to hypothesize that its efficacy may be linked to its lung targeting. Below, we test Abraxane’ therapeutic efficacy against lung metastases in wild type mice, where it targets the lungs, and in C3 knockout mice, where it does not target the lungs (Figure 1e).

B16-F10 melanoma cells transfected with firefly luciferase (fLuc) and tdTomato reporter transgenes were administered intravenously in wild type and C3 knockout mice. Over 18 days, tumor-associated luciferase activity was measured twice weekly by luminescence imaging after intraperitoneal luciferin. Luminescence was highly concentrated in the lungs, indicating lung metastases seeding via B16-F10 embolism in all treatment groups. The mice were treated with either 5 mg/kg Abraxane or phosphate buffered saline sham intravenously every other day for the first 10 days after injection of melanoma cells.

Melanoma reporter luminescence in the lungs accumulated over three weeks in all groups. Relative to sham-treated control, tumor-associated luminescence trends towards reduced signal at 18 days in wild type mice treated with Abraxane, agreeing with published results showing Abraxane reduces tumor growth in similar models (Figure 5a-c)(*36*). When Abraxane’s lung uptake is ablated in C3 knockout mice, Abraxane fails to impede growth of B16-F10 tumors arising from lung metastases (Figure 5d-f). We observe no significant effects on survival or body weight for Abraxane treatment in either genotype (Supplementary Figure 26).

**Figure 5.**
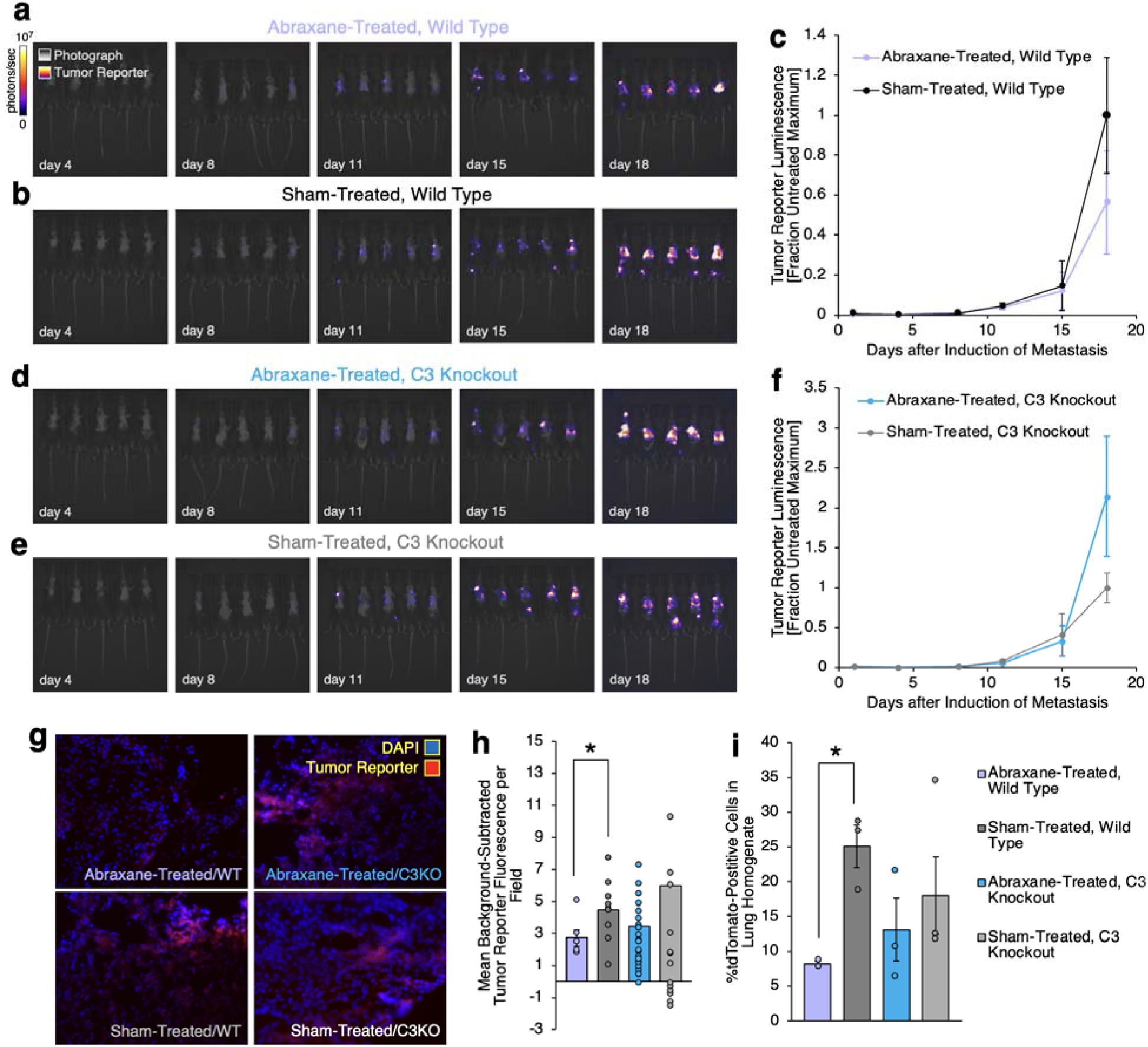
**Intravenous Abraxane effects on lung metastases in wild type vs. complement C3 knockout lungs**. (a-b) Imaging of B16-F10 melanoma cell reporter luminescence in mice at 4, 8, 11, 15, and 18 days after intravenous injection of 4x10^5^ cells per mouse. a: Mice were treated with 5 mg/kg IV Abraxane bolus at 0, 2, 4, 6, and 8 days after IV B16-F10 injection. b: Mice received intravenous PBS sham on the same schedule as Abraxane boluses. (c) Summary data averaging B16-F10 luminescence (photons/second flux) in regions of interest around the lungs in images depicted in (a-b). Regions of interest with identical dimensions were used for each mouse at each time point. n=5 mice per group. (d-e) Images as in (a-b), but in complement C3 knockout mice, rather than wild type mice. (f) Summary data as in (c) for imaging data in (d-e). n=5 mice per group. (g-h) Representative images depicting B16-F10 reporter fluorescence in lung tissue 21 days after IV B16-F10 injection (g) and quantification of mean fluorescence in imaging fields at 20X magnification (. Mice treated as described in (a-b). Sham-treated wild type: n=9 imaging fields over 3 mice, Abraxane-treated wild type: n=6 imaging fields over 2 mice, sham-treated C3 knockout: n=15 imaging fields over 5 mice, Abraxane-treated C3 knockout: n=23 imaging fields over 5 mice. *p=0.05 by unpaired t-test. (i) Quantification of of tdTomato reporter-positive cells in flow cytometry data indicating tumor reporter fluorescence as a function of Abraxane treatment in single cell suspensions prepared from lungs from mice treated as in (a-b). *p=0.05 by unpaired t-test.s

Post-mortem, tumor load in the lungs was assessed via tdTomato reporter fluorescence in histology and flow cytometry data. Histology confirmed the presence of tdTomato fluorescence in the lungs from all treatment groups. Evaluating background-corrected fluorescence per imaging field in Abraxane- vs. sham-treated lungs, we find significantly reduced reporter fluorescence in Abraxane-treated wild type lungs. In C3 knockout mice, Abraxane exerts no significant effect on tumor fluorescence (Figure 5g-h, Supplementary Figures 27-28).

In flow cytometry data, frequency of tdTomato-positive cells was taken to represent frequency of B16-F10 cancer cells in single cell suspensions prepared from affected lungs (Supplementary Figure 29). The frequency of tdTomato-positive cells in the lungs is reduced by Abraxane treatment in wild type mice, but not in C3 knockout mice (Figure 5i). Collectively, this portion of the results shows that Abraxane does not reduce growth of tumors arising from lung metastases in C3 knockout mice, where Abraxane’s lung uptake is ablated.

## Discussion

At first glance, it may seem that our findings diverge from previous assessments of Abraxane. Prior biodistribution studies have described unremarkable accumulation of paclitaxel in the lungs following intravenous Abraxane administration(*14, 28*). However, these studies trace drug liberated from the albumin component of Abraxane, using extraction techniques that may struggle to distinguish the nanoparticle form of the drug(*37*). Our study instead focuses on the albumin component of Abraxane. In effect, we have considered how paclitaxel affects albumin, rather than the other way around. We have used a tracing strategy that is more likely to reflect the fate of the intact nanoparticle, rather than drug eluted from the nanoparticle. Tracing albumin, the carrier component of Abraxane, led us to evidence for uptake of Abraxane in the lungs, in agreement with another recent study finding the same with fluorophore-labeled Abraxane(*17*).

Prior studies have demonstrated delivery of nanoparticles with properties resembling those of Abraxane to marginated neutrophils in inflamed lungs (*18, 38, 39*). Our studies of Abraxane in acute systemic and pulmonary inflammation show that Abraxane’s lung uptake can be elevated following inflammation. That finding led us to show that neutrophils, a cell type that localizes to the pulmonary vasculature in an inflammation-dependent manner(*40*), take up Abraxane. This finding is consistent with the prior literature studying nanoparticles resembling Abraxane.

We’ve demonstrated that neutrophil-driven Abraxane lung uptake depends on the complement pathway. Other studies of neutrophil/lung-tropic nanoparticles show that C3b, a fragment of the complement protein C3, can coat nanoparticle surfaces(*20, 41*), leading to recognition by receptors for C3b on neutrophils and other phagocytes. Here, C3 knockout eliminates neutrophil/lung tropism of Abraxane, implying a similar mechanism. For nanoparticles with lung tropism and other nanomedicines, the complement pathway is a source of acute side effects. Acute complement pathway toxicity is largely mediated by the complement protein fragments C3a and C5a(*42*). These anaphylatoxins are released into plasma as a byproduct of C3b adhesion to nanoparticle surfaces. However, acute complement responses have not been documented with Abraxane. Since C3a and C5a are rapidly cleared from circulation(*43, 44*), some nanoparticles that accumulate C3b on their surfaces may not provoke an anaphylactic response. Indeed, our measurements show that C3a and C5a levels are not significantly elevated in plasma following Abraxane bolus dosing (Supplementary Figure 30).

Even in the absence of a robust C3a and C5a response, C3b on nanoparticle surfaces can engage with neutrophils and other phagocytic cells(*45*). Phagocytosis of nanoparticles can provoke neutrophil activation and adhesion along the vascular surface. Neutrophil activation leads to release of granules containing proinflammatory and prothrombotic factors(*40*). Activated neutrophil surfaces, presented along the vasculature following adhesion, attract platelets and additional leukocytes(*40, 46*). Clinical data show Abraxane causing neutropenia(*47, 48*), which can be exacerbated when neutrophils leave circulation to adhere along the vasculature. Our data show that Abraxane provokes lung-specific elevation of IL-6 and thrombi, consistent with these neutrophil-driven mechanisms.

Complement-mediated Abraxane interactions with neutrophils should be understood as a potential source of side effects(*41, 42, 45*). However, the side effects we observed were unique to lungs with pre-existing inflammation, induced by intravenous LPS. This implies that Abraxane requires a prior agonist to induce effects on IL-6 and coagulation. No such effects have been noted in clinical data, but our findings might be considered in the evaluation of risk when Abraxane patients suffer from acute inflammatory disorders.

Homing of Abraxane to the lungs is itself a newly observed effect. We show that this effect may be entwined with Abraxane’s therapeutic efficacy. When Abraxane’s lung uptake is inhibited by complement knockout, Abraxane is not effective against lung tumors in mice. Our experiments directly focus on metastases in the lungs, but the significance of the lung vasculature as a landing site for metastases in general(*49*) might explain the success of Abraxane in metastatic pancreatic and breast cancer and offer hope for expanded use of Abraxane in other metastatic cancers.

Abraxane’s interactions with neutrophils explain its lung uptake and may also point to new treatment modalities. Activated neutrophils colocalize with circulating cancer cells at the onset of metastasis and tumor-associated neutrophils could be a good carrier cell for surveilling prevention of metastasis(*24, 25*). Strategies for targeting neutrophils may be pursued to replicate Abraxane’s success. Provocation of apoptosis in neutrophils colocalized with tumor cells has been proposed to enhance immune responses against cancer. Indeed, prior work has shown promising results treating mouse model tumors with neutrophils provoked to apoptosis by *ex vivo* loading with Abraxane(*26*). Of note, in metastatic pancreatic cancer patients where Abraxane and Gemcitabine cause more profound neutropenia, survival is improved(*47*).

In summary, we have presented a new mechanism by which to understand Abraxane. Abraxane has variously been proposed to improve pharmacokinetics of paclitaxel, enhance retention in the leaky vasculature of solid tumors, or engage with albumin receptors in and around the tumor vasculature(*2, 3, 12, 17*). Our study doesn’t exclude these mechanisms but adds a new one: Physicochemical targeting to neutrophils. Nanomedicine has used engineering of nanoparticle chemistry and structure to achieve tropisms for different organs and cell types(*21–23*). Along these lines, our prior work has characterized motifs in nanoparticle structure that predict uptake in pulmonary neutrophils(*18*). The established structure of Abraxane closely matches these motifs and we show here that Abraxane indeed targets pulmonary neutrophils via mechanisms consistent with that physicochemical tropism.

## Methods

### Nanoparticle and protein radiolabeling and fluorescence tagging

Abraxane was obtained from pharmacy stock. Lyophilized Abraxane was stored at -80°C prior to gentle resuspension at 5 mg/mL drug concentration in phosphate buffered saline without calcium or magnesium. The suspensions were diluted to 0.05 mg/mL concentration in PBS for assessment of size and polydispersity with dynamic light scattering (DLS, Malvern Zetasizer). The suspensions were diluted to 0.5 µg/mL for assessment of size and particle concentration with nanoparticle tracking analysis (NTA, Malvern NanoSight).

For biodistribution studies, Abraxane was labeled with ^125^I. 300 µL of 0.5 mg/mL Iodogen (Perkin-Elmer) was dried in a borosilicate tube under nitrogen gas. 100-200 µL of 5 mg/mL Abraxane suspension was added to the tube immediately before Na^125^I at 25 µCi per 100 µg of Abraxane. The Abraxane-sodium iodide solution was incubated for 5 minutes at room temperature after covering with parafilm. Both incubation and Na^125^I solution preparations were performed in a ventilated chemical hood. Radiolabeled Abraxane was separated from excess Na^125^I with 7 kDa cutoff gel filtration columns (Zeba). Radioactivity in the Abraxane sample and in the columns was quantified with a gamma counter (Wizard). Abraxane was passed through successive columns until >95% of ^125^I signal was associated with the nanoparticles, rather than free in solution. Association of ^125^I with the nanoparticles was confirmed with thin film chromatography (TLC). 1 µL of labeled Abraxane was applied near the end of a silica gel strip and that end of the strip was submerged in 10 mM EDTA at a depth that did not cover the point at which the Abraxane was applied. The mobile phase EDTA solution was allowed to advance along the strip until the solvent front was ∼1 cm from the opposite end. ^125^I signal within ∼1cm of the initial sample location was taken to quantify ^125^I retained in Abraxane and signal from the rest of the strip was taken to quantify unbound ^125^I. After iodination, Abraxane size, polydispersity, and concentration were assessed with DLS and NTA as above.

For tracing in flow cytometry and histology experiments, Abraxane was labeled with NHS ester Alexa Fluor 647 (ThermoFisher). NHS ester Alexa Fluor 647 was prepared as an 8 mM solution in dry DMSO and added to Abraxane suspensions at 5 uL per 10 mg nanoparticles. The Abraxane-fluorophore mixture was gently rotated for two hours at 4°C. Excess fluorophore was separated from the Abraxane by centrifugation against 10 kDa molecular weight cutoff centrifugal filtration units (Amicon). Labeled Abraxane was rinsed and filtered four times and lack of fluorophore in the final filtrate/conjugation of fluorophore to Abraxane was confirmed by optical density of the samples at 647 nm (Nanodrop spectrophotometry). Size, polydispersity, and concentration of fluorophore-labeled Abraxane was assessed with DLS and NTA.

### Biodistributions in mouse models

Abraxane and albumin were labeled with iodine radioisotopes as described above. ∼0.2 µCi of radioactive Abraxane was reserved for each mouse, to be doped into sufficient unlabeled Abraxane to provide a 1 mg/kg or 5 mg/kg intravenous bolus drug dose. After anesthetizing mice with 3% isoflurane, Abraxane or equivalent dose of albumin control was injected via the jugular vein over 1 minute. By gamma counter, we determined the amount of radioactivity loaded into the syringes prior to injection and the amount of radioactivity remaining in the syringes after injection, recording the injected dose as the difference between loaded and residual radioactivity. 30 minutes after injection, blood was drawn from the inferior vena cava, followed by severing the vena cava to terminally exsanguinate. Organs were excised, rinsed in saline or PBS without calcium or magnesium, blotted dry, and placed in culture tubes for measurement of isotope-specific radioactivity in a gamma counter (Perkin-Elmer). Radioactivity measured in each organ and in 100 µL blood samples was compared to injected radioactivity to determine the fraction of injected dose distributing to different organs.

Biodistributions were evaluated in naïve mice, mice treated with intravenous LPS, and mice treated with inhaled H1N1 influenza. For intravenous LPS, LPS from E. coli strain B4 was injected at a 2 mg/kg dose via the retro-orbital cavity after anesthetizing with 3% isoflurane. Mice were dosed with Abraxane 5 hours after LPS, as described above. For H1N1 infection, PR8-GP33 H1N1 influenza at a dose of 0.3 LD_50_ (as determined in previous work by the Morrisey lab) was administered to mice intranasally in 30 µL of PBS. 14 days after infection, mice were dosed with Abraxane as described above.

### Complete blood count analysis

From naïve and intravenous LPS-treated mice receiving a 5 mg/kg intravenous bolus of Abraxane, 100 µL of drawn blood was added to an EDTA-coated tube and evaluated with a three-part hematology analyzer (VetScan HM5). Identical analyses were performed for blood from naïve and intravenous LPS-treated mice receiving no Abraxane.

### Ektacytometry

In blood samples from naïve and intravenous LPS-treated mice receiving 5 mg/kg intravenous Abraxane, we evaluated red blood cell mechanical properties via ektacytometry (RheoScan-D, RheoMeditech). 5 µL of whole blood was dispersed in 695 µL of 5.5% w/v 360 kDa polyvinylpyrrolidone in PBS to create homogeneous suspensions. Using RheoScan-D cassettes, cells in the suspensions were subjected to ∼0-20 Pa shear. Diffraction patterns generated by red blood cells in the suspensions were recorded. RheoScan-D software fit the diffraction patterns to ellipses, recording values for length of the long and short axes of each ellipse (*L* and *S*, respectively) and the shear stress at which the corresponding ellipsoidal diffraction pattern was generated (*Tau*). RBC elongation indices were defined as *EI* = (*L* – *S*)/(*L* + *S*). Plots of *EI* vs. *Tau* were used to characterize the response of RBCs in the blood samples to shear stress.

### Effects of Abraxane in nebulized LPS-induced acute lung injury

A 5 mg/mL solution of B4 LPS in PBS was prepared and 5 mL of the LPS solution was aerosolized in a nebulizer (NEB-MED H, Braintree Scientific) connected to an exposure chamber with separate compartments for individual mice (MPC-3 AERO, Braintree Scientific). To improve survival over the course of the injury, mice received 1 mL of 37°C sterile saline intraperitoneally immediately before exposure to LPS. Mice were exposed to LPS over ∼20 minutes, during which time all liquid in the nebulizer was aerosolized. Using methods described above in *Biodistributions in mouse models*, 5 mg/kg intravenous boluses of Abraxane or intravenous PBS sham doses were administered 2 hours after LPS exposure. Mouse body weights were determined before LPS exposure and 24 hours after LPS exposure.

After determining body weights at 24 hours after LPS exposure, bronchoalveolar lavage fluid was obtained. Briefly, mice were anesthetized with intramuscular ketamine-xylazine (10 mg/kg ketamine, 100 mg/kg xylazine) and tracheostomies were performed with 22-gauge catheters. After euthanizing by severing the inferior vena cava, 0.8 mL of cold 0.5 mM EDTA in PBS was injected into the lungs over 1 minute via the 22-gauge tracheostomy catheter. The buffer was aspirated from the lungs over an additional 1 minute. Two additional 0.8 mL injections/aspirations were performed to further rinse the bronchoalveolar space. Recovered bronchoalveolar lavage fluid was centrifuged at 300xg for 4 minutes. The resultant supernatants were collected and stored at -80°C prior to analysis of protein concentration (Bio-Rad DC Protein Assay). Cell pellets were fixed by resuspending in 333 µL of 1.6% PFA in PBS and incubating for 10 minutes in the dark at room temperature. 1 mL of 0.5 mM EDTA in PBS was added after fixation, followed by centrifugation at 400xg for 3 minutes, removal of the resulting supernatant, and resuspension of the cell pellet in 1 mL of PBS containing 2% fetal calf serum and 1 mM EDTA. These suspensions were centrifuged 400xg for 3 minutes and, after removal of the supernatants, the cell pellets were resuspended in 100 µL of the same buffer containing a 1:1000 dilution of APC-labeled anti-CD45. The samples were stained for 30 minutes at room temperature in the dark, then 1 mL of PBS containing 2% fetal calf serum and 1 mM EDTA was added. The suspensions were centrifuged at 400xg for 3 minutes, the supernatant was discarded and the cells were finally resuspended in 900 µL of the same buffer prior to flow cytometry analysis. After gating out debris and doublets according to the strategy depicted in Supplementary Figure 8, anti-CD45 stained samples were compared to unstained samples to delineate and count leukocytes in the bronchoalveolar lavage fluid.

### Mouse lungs histology

30 minutes after intravenous administration of a 5 mg/kg intravenous bolus of Abraxane or an equivalent volume of a PBS sham in naïve or intravenous LPS-treated mice (see *Biodistributions in mouse models* above), the inferior vena cava was severed and the right ventricle was perfused with ∼10 mL of cold PBS. The trachea was cannulated with a 22-gauge catheter (see *Effects of Abraxane in nebulized LPS-induced acute lung injury* above) and the lungs were inflated and fixed with 10% formalin buffered to neutral pH. The fixed lungs were embedded in paraffin and sectioned at 5 µm thickness. Lung sections were stained via Masson’s trichrome technique and imaged by the Pathology Core Laboratory of the Children’s Hospital of Philadelphia. From each of the four treatment groups, four images at 10x magnification and four images at 20x magnification were selected by random number generator. Of the 32 selected images, each was assigned a label for blinded scoring by a second round of random number generation. Two blinded scorers were tasked with counting the number of thrombi in each randomly labeled field of view.

To image distribution of fluorescent Abraxane in mouse lungs, 30 minutes after intravenous administration of a 5 mg/kg intravenous bolus of Alexa Fluor 647-labeled Abraxane (see *Nanoparticle and protein radiolabeling and fluorescence tagging* above) in naïve or intravenous LPS-treated mice (see *Biodistributions in mouse models* above), mice were exsanguinated, lungs were perfused, and the tracheae were canulated as described above. Via the trachea, lungs were filled with M1 embedding matrix and immediately frozen in liquid nitrogen. The frozen lungs were sectioned to 15 µm thickness at -20°C with a Leica CM 1950 cryostat. Section on glass slides were fixed by incubating 15 minutes in 2% paraformaldehyde at room temperature. After blocking with 1% bovine serum albumin and 3% triton in PBS for one hour at room temperature, staining antibodies were added to the slides at a 10 µg/mL working concentration in the blocking buffer. Alexa Fluor 594 conjugated antibody stained for CD31 and FITC-conjugated antibody stained for CD45. After staining, slides were rinsed with blocking buffer and mounting media containing DAPI was used to fix the sections under cover glass prior to confocal imaging. Confocal images were analyzed in ImageJ.

### Mouse lungs flow cytometry

30 minutes after intravenous administration of a 5 mg/kg intravenous bolus of Alexa Fluor 647-labeled Abraxane (see *Nanoparticle and protein radiolabeling and fluorescence tagging* above) in naïve or intravenous LPS-treated mice (see *Biodistributions in mouse models* above), lungs were isolated to prepare single-cell suspensions. As in *Effects of Abraxane in nebulized LPS-induced acute lung injury* above, mice were anesthetized with intramuscular ketamine-xylazine and their tracheas were cannulated with a 22-gauge catheter. The catheter was secured in place by suture around the trachea, mice were euthanized by severing the inferior vena cava, and the lungs were perfused by instillation of ∼10 mL cold PBS through the right ventricle. 1 mL of cold PBS with 5 U/mL dispase, 2.5 mg/mL collagenase I, and 1 mg/mL of DNAse I was injected into the tracheal catheter, then a suture was tightly tied around the trachea as the catheter was removed. The lungs, trachea, and heart were removed by thoracotomy.

In a petri dish on ice, lung tissue was isolated from the heart and trachea, and the lungs were disaggregated into slurries by triturating. The slurries were aspirated into fresh tubes, then diluted with an additional 2 mL of dispase/collagenase/DNAse solution. The suspensions were incubated for 45 minutes at 37°C, vortexing every 10 minutes. The digested suspensions were then passed through 100 µm cutoff cell strainers into 50 mL conical tubes and the filtered suspensions were centrifuged at 400xg for 5 minutes. The supernatant was discarded and the pellet was resuspended in 10 mL cold ACK lysing buffer. After 10 minutes incubation on ice, those suspensions were passed through 40 µm cutoff cell strainers into new 50 mL conical tubes, then centrifuged at 400xg for 5 minutes once more. The supernatant was discarded and the pellet was rinsed with 10 mL of PBS with 2% fetal calf serum and 1 mM EDTA. Following another 400xg/5-minute centrifugation, the supernatant was discarded and the cells were resuspended in 2% paraformaldehyde in 1 mL of PBS with 2% fetal calf serum and 1 mM EDTA. That suspension was incubated in the dark for 10 minutes at room temperature, followed by centrifugation at 400xg for 5 minutes, removal of the resulting supernatant, and re-suspension in 1 mL PBS with 2% fetal calf serum and 1 mM EDTA.

100 µL aliquots of the fixed cell suspensions were stained by pelleting at 400xg for 5 minutes then resuspending in solutions of labeled antibodies in PBS with 2% fetal calf serum and 1 mM EDTA. After incubation for 20 minutes in the dark at room temperature, the 100 µL staining suspensions were diluted with 1 mL of PBS with 2% fetal calf serum and 1 mM EDTA and centrifuged at 400xg for 5 minutes. Supernatants were discarded and the stained pellets were resuspended in 200 µL of PBS with 2% fetal calf serum and 1 mM EDTA.

Data for stained cell aliquots were obtained on a BD Accuri C6 Plus flow cytometer immediately after resuspension. Staining antibodies were PE-labeled anti-CD45 (BD Biosciences, 1:500 dilution during staining), FITC-labeled anti-CD31 (Invitrogen, 1:150 dilution), and PerCP/Cy5.5-labeled anti-Ly6G (BD Biosciences, 1:150 dilution). Single cell suspensions were also prepared from naïve and intravenous LPS-treated mice receiving no labeled Abraxane, to provide unstained controls. The unstained controls and single-color stained controls provided data to automatically create compensation matrices for naïve and LPS-injured groups in FlowJo software. All other gating relied on comparison of unstained and single-stained samples.

### IL-6 and MIP-2 measurements

Concentrations of the cytokine IL-6 and the chemokine MIP-2 were measured in plasma and in homogenized lung and liver tissue. Lung and liver tissue were homogenized in 1 mL of PBS containing protease inhibitor cocktail (Sigma) at 1x concentration. 50 mM Tris-HCl with 2 mM EDTA at pH 7.4 was added to homogenized tissue or plasma at 100 mg of tissue per 900 µL buffer and the mixtures were incubated for one hour at 4°C. The lysates were centrifuged at 16000xg for 10 minutes at 4°C and the supernatants were recovered. IL-6 and MIP-2 concentrations in the supernatants were determined using ELISA kits (DuoSet, R&D Systems) according to manufacturer’s instructions.

### Human lungs experiments

Human lungs were obtained after organ harvest from transplant donors whose lungs were in advance deemed unsuitable for transplantation due to radiographic evidence of alveolar filling and low PaO_2_ to FiO_2_ (P/F) ratios. The lungs were harvested by an organ procurement team and kept at 4°C until the experiment, which was done within 24 hours of organ harvest. The lungs were inflated with low pressure oxygen and the main bronchus was clamped. Oxygen flow was maintained at ∼0.8 L/min to maintain gentle inflation. Pulmonary artery subsegmental branches were endovascularly cannulated, then tested for retrograde flow by perfusing for 5 minutes with Steen solution containing 3% BSA doped with green tissue dye and then sealed to the artery opening with tissue glue composed of 30% bovine serum albumin (BSA) mixed at 1:1 with glutaraldehyde, thus preventing retrograde efflux of solutions injected into the artery branch. The lungs were perfused with 3% BSA in PBS at 25 cm H_2_O pressure. The pulmonary veins through which efflux of perfusate emerged were noted, allowing collection of solutions after passage through the lungs. A 2 mL mixture of ^125^I-labeled Abraxane, ^125^I-labeled Abraxane and ^131^I-labeled human albumin, or Alexa Fluor 647-labeled Abraxane was injected through the arterial catheter. ∼100mL of 3% BSA in PBS was passed through the same catheter to rinse unbound nanoparticles or protein. A solution of green tissue dye was subsequently injected through the same catheter. The cannulated lung lobe was dissected into ∼1 g segments, which were evaluated for density of tissue dye staining. For radiolabel tracing experiments, segments were weighed, divided into ‘high’, ‘medium’, ‘low’, and ‘null’ levels of dye staining, and measured for ^131^I and ^125^I signal in a gamma counter. For flow cytometry and histology experiments ∼0.25 g segments with high levels of tissue dye staining or similar tissue segments from non-perfused portions of the lungs were reserved for further tissue processing (flow cytometry) or frozen in liquid nitrogen for histology.

To prepare human lung segments for flow cytometry, tissue was digested with a solution of 442.5 U/mL Collagenase Type I (Life Tech), 5 U/mL Dispase (Fisher), and 9.9 U/mL DNase1 in PBS without calcium and magnesium. Segments of ∼0.5 g mass were triturated with surgical scissors and the resulting macerated sample was added to 6 mL of the digestion solution warmed to 37°C. The tissue/digest buffer was incubated at 37°C for 35 minutes, vortexing at high speed every 10 minutes. The resulting suspensions were filtered through 70 µm cutoff cell strainers and the strainers were washed with 10 mL PBS. The filtered suspensions were centrifuged at 350xg for 5 minutes. The resulting supernatants were discarded and the pellets were gently resuspended in 10 mL ACK lysis buffer to destroy red blood cells in the suspensions. After 10 minutes incubation at room temperature, the ACK suspensions were centrifuged at 350xg for 5 minutes and supernatants were discarded. The pellets were washed with 10 mL of FACS buffer (PBS with 0.2% EDTA and 1% FBS) and centrifuged at 350xg for 5 minutes. Supernatants were discarded, pellets were resuspended in 5 mL FACS buffer, and the resulting suspensions were passed through 40 µm cutoff cell strainers, with 10-15 mL FACS buffer used to wash remaining cells through the strainers. After centrifuging at 350xg for 5 minutes and discarding supernatants, the final cell pellets were resuspended in 5 mL FACS buffer and cell concentrations were determined. Cell concentrations were adjusted to 5 million cells per mL and incubated with Fc blocking agent (Biolegend) at 1:40 concentration for 15 minutes at 4°C. After washing away blocking agent by centrifugation, cells were stained in 200 µL aliquots at 5 million cells per mL. Two staining panels were employed with all staining antibodies diluted to 1:200 concentration: 1) Antibodies against CD45 (FITC-conjugated, Biolegend), CD206 (PE-Cy7-conjugated, Thermofisher), CD15 (BV421-conjugated, Biolegend), and CD169 (PE-conjugated, Biolegend); 2) Antibodies against CD45 (FITC-conjugated, Biolegend), CD31 (PE-conjugated, BD), and EpCAM (PerCP-Cy5.5-conjugated, Biolegend). After staining for 30 minutes at 4°C, cells were pelleted at 350xg for 5 minutes and washed with FACS buffer. Stained cells were fixed with 4% PFA for 30 minutes at 4°C, washed as above, then analyzed with a BS LSRFortessa flow cytometer. Data was analyzed with FlowJo software according to the gating strategy depicted in Supplementary Figure 20.

To prepare lung segments for histology, tissue was submerged in optimal cutting temperature gel and frozen in liquid nitrogen prior to storage at -80°C. Frozen tissue was sectioned at 15 µm thickness and mounted on glass slides at -20°C using a Leica CM 1950 cryostat. Sections were fixed in 2% paraformaldehyde at room temperature for 15 minutes. Slides were incubated in 1% bovine serum albumin and 3% triton in PBS without calcium or magnesium for one hour at room temperature. Staining antibodies were added to the slides at ∼10µg/mL working concentration in two stain sets: 1) Antibodies against CD45 (FITC-conjugated, Biolegend) and CD15 (BV421-conjugated, Biolegend); 2) DAPI and antibody against CD45 (FITC-conjugated, Biolegend). Slides were stained overnight at 4°C. After gently washing with 3% triton in PBS, slides were mounted under cover glass and imaged with an LSM980 confocal microscope (Zeiss). Images were prepared for figures in FIJI.

### Abraxane treatment in B16-F10 lung metastasis model

Luciferase/tdTomato dual reporter B16/F10 melanoma cells were purchased from ALSTEM. Per manufacturer’s instructions, 1x10^6^ cells were thawed at 37°C, added to 10 mL of culture medium (Dulbecco’s Modified Eagle Medium/DMEM with 10% fetal bovine serum/FBS and 1% antibiotic/antimycotic), centrifuged at 200xg at room temperature, resuspended in 1 mL of culture medium, and added to 5 mL of culture medium in a T25 flask at 37°C in an incubator maintained at 5% CO_2_. Cells were grown to 85% confluence before transferring to T-175 flasks. At each passage, cells were dissociated with Gibco TrypLE Express and, prior to re-plating, a portion of the cell pellet was reserved to confirm maintenance of tdTomato signal with widefield fluorescence microscopy. Cells were grown through five passages prior to use in mice.

Cells were dissociated and rinsed prior to suspension in cold PBS at a concentration of 2.5x10^6^ cells per mL. Following procedures in Bhattarai et al.(*36*), 2.5x10^5^ B16/F10 melanoma cells were injected intravenously in each of 10 C57BL/6 mice and 10 C3 knockout mice (Jackson Laboratories) anesthetized via 3% isoflurane. Among the C57BL/6 mice and the C3 knockout mice, five mice from each group were randomly selected to receive five doses of intravenous Abraxane at 5 mg/kg drug dose on alternating days. All other mice received intravenous PBS on an identical schedule.

Mice were imaged at 1, 4, 8, 11, 15, and 18 days over the course of tumor growth to assess B16/F10-associated luciferase-induced luminescence in the lungs. To prepare for in vivo imaging system (IVIS, Revvity Spectrum) experiments, mice were put under anesthesia using 3% isoflurane, depilated over their thoraxes, and intraperitoneally injected with 100µL of 30 mg/mL D-luciferin sodium salt (Regis Technologies). Anesthetized mice were positioned in supine orientation to view the thorax and imaged for chemiluminescence every minute with automatically determined exposure time. 10-14 images were obtained to identify peak signal intensity. Revvity LivingImage software (version 4.8.2) was used to analyze images, identifying reporter luminescence intensity in regions of interest focused on the chest and with identical geometry for each mouse.

Lungs were collected from mice either after failure to survive the 22-day time course of the experiment or after terminal exsanguination. The left or right lungs from each mouse were randomly selected either for preservation for fluorescence immunohistochemistry or for processing for analysis by flow cytometry. For immunohistochemistry, lung lobes were embedded in optimal cutting temperature gel, frozen in liquid nitrogen, and preserved at -80°C. As with methods for human lung segments, the lobes were sectioned at 15 µm thickness at -20°C using a Leica CM 1950 cryostat. Lung sections were mounted under coverglass with DAPI mounting media and imaged to detect tdTomato fluorescence and DAPI signal. 3-5 regions of interest were selected from each slide to represent regions of the lung lobes with maximum tdTomato signal. Mean tdTomato fluorescence in each region of interest was determined with FIJI ImageJ software, after omitting portions of the image identified by DAPI staining as outside of the tissue. For flow cytometry analysis, lungs were processed as described in *Mouse lungs flow cytometry* above, but with no staining applied to the cell suspensions. Lungs from a C57BL/6 mouse receiving no B16/F10 melanoma cells and no Abraxane were prepared identically to provide a cell suspension with no tdTomato signal. The single cell suspensions were analyzed with a Cytek Guava flow cytometer to quantify tdTomato fluorescence intensity. tdTomato fluorescence intensity was represented in histograms and frequency of tdTomato-positive cells was counted using FlowJo version 10.10.0.

### Statistical analysis

All statements of statistical significance are noted in figure legends, alongside descriptions of applied statistical analyses and numbers of replicates in the relevant experiments. Statistical analyses were performed in Graphpad Prism, version 10.

## Acknowledgments

We thank the Penn Cytomics and Cell Sorting Core (RRID: SCR_022376) and Cell & Developmental Biology Microscopy Core (RRID: SCR_022373) at the University of Pennsylvania. Penn Cytomics is partially supported by Abramson Cancer Center NCI Grant (P30 016520).

## Funding

NIH R01-HL157189 (V.R.M.)

NIH R01-NS131279 (O.A.M-C.)

NIH R01-HL153510 (J.S.B.)

NIH R01-HL160694 (J.S.B.)

NIH R01-HL164594 (J.S.B.)

Pennsylvania Department of Health Research Formula Fund (V.R.M.)

National Center for Advancing Translational Sciences of the National Institutes of Health award UL1TR001878 (J.W.M.)

## Author Contributions

J.S.B, O.A.M-C., and J.W.M. conceptualized the study, designed the experiments, interpreted the data, and wrote the manuscript. A.M., S.J., M.E.Z., N.M., Y.W., J.N., T.B., P.P., M.H.Z., J.W., C.T.S., L.S.C., Z.W., L.F., O.A.M-C., and J.W.M. performed experiments and processed data. A.M., S.J., N.M., T.B., O.A.M-C., and J.W.M. performed experiments in models of lung metastasis. Y.W., J.N., and J.W.M. obtained histology data. A.M., J.W., L.S., and J.W.M. performed experiments in human lungs. M.E.Z. and J.W.M. obtained flow cytometry data. A.M., S.J., M.E.Z., P.P., M.H.Z., C.T.S., L.S.C., Z.W., O.A.M-C., and J.W.M performed experiments tracing biodistributions and side effects in mice. All authors approved the manuscript.

## Data and Materials Availability

All data are available in the main text or the supplementary materials. Unprocessed tabulated forms of the data are available on reasonable request to the corresponding authors.

