## Supplemental Material for "Nanoparticle Albumin-Bound Paclitaxel Targets Pulmonary Neutrophils"

### Supplementary Figures

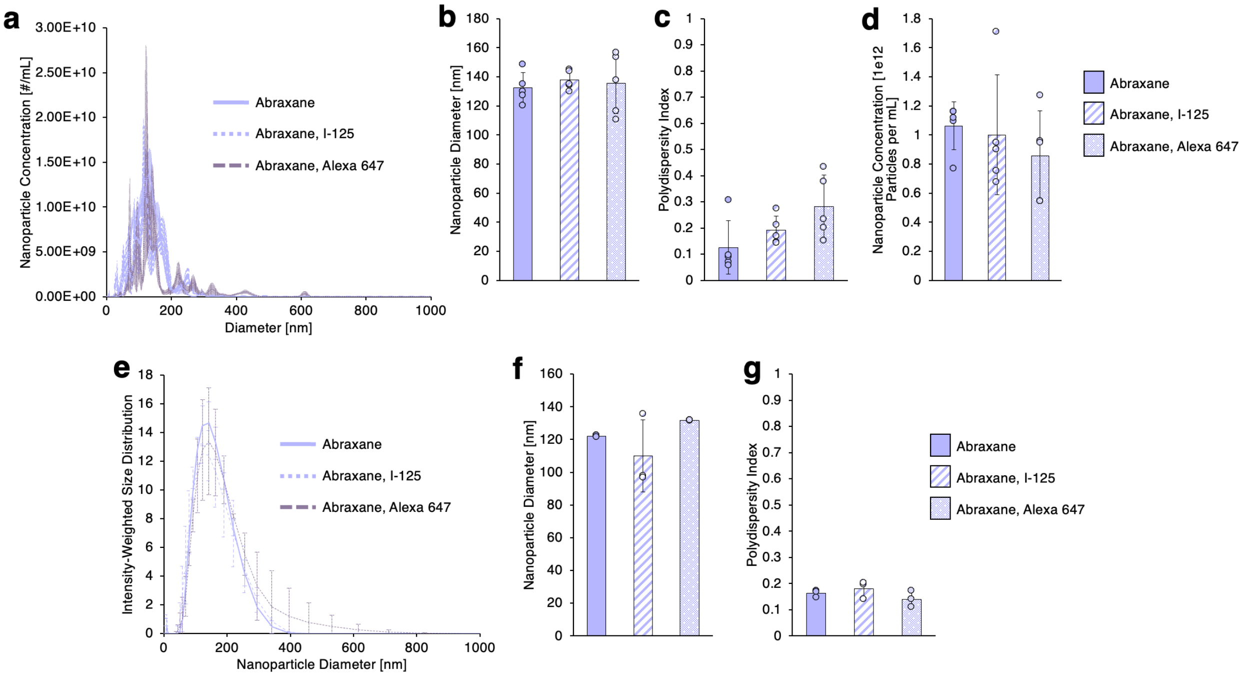


Supplementary Figure 1. Characterization of Abraxane, radiolabeled Abraxane, and Alexa Fluor 647 labeled Abraxane. (a-d) Nanoparticle tracking analysis data. (a) Distributions showing size-resolved concentrations of 1:10000 dilutions of Abraxane, radiolabeled Abraxane, or fluorophore-labeled Abraxane, from recommended reconstitution concentration. (b) Abraxane diameters derived from size distributions in (a), showing no significant change in diameter due to labeling with $^{125}$I or fluorophore. (c) Statistical polydispersity indices of Abraxane, as calculated from size distributions in (a). (d) Abraxane concentration measurements, showing no loss of material due to labeling. (e-f) Dynamic light scattering data. (e) Intensity-weighted size distributions for Abraxane, radiolabeled Abraxane, or fluorophore-labeled Abraxane. (f) Nanoparticle diameters derived from the size distributions in (e), showing that Abraxane's diameter is not significantly changed by labeling with $^{125}$I or Alexa Fluor 647. (g) Polydispersity indices derived from the size distributions in (e).

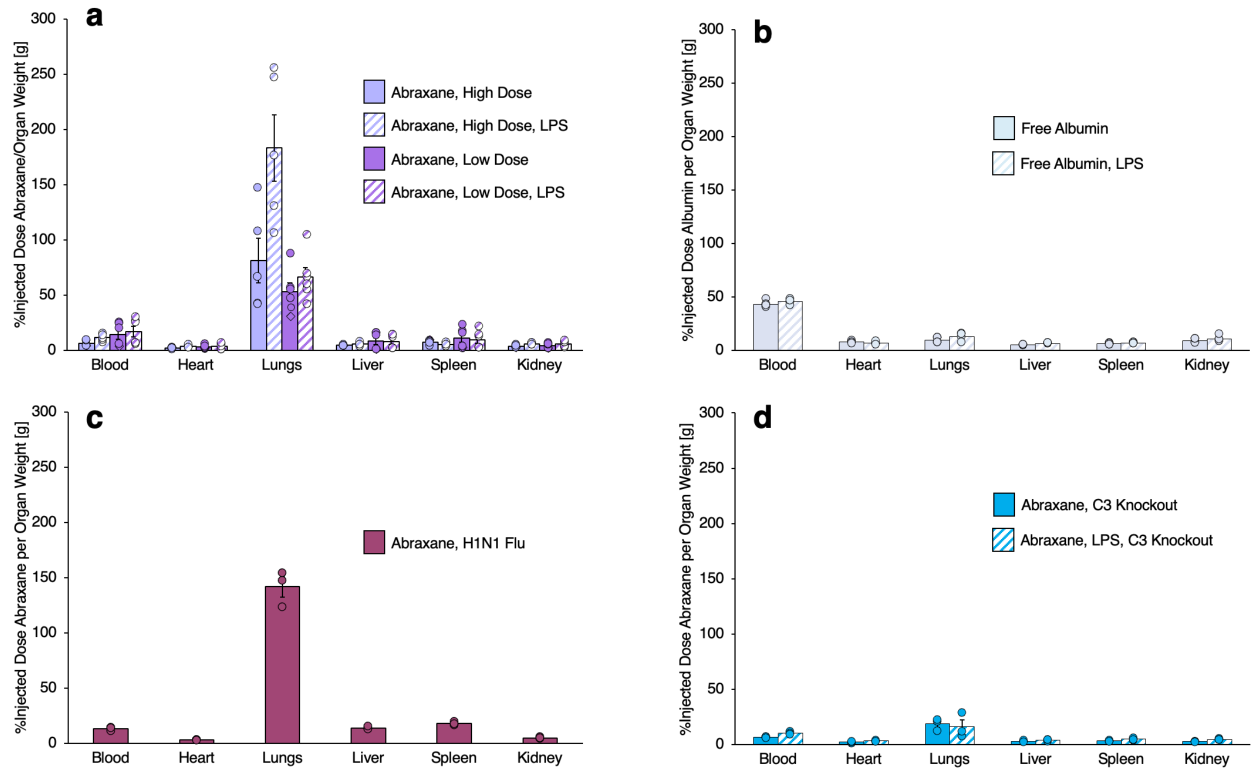


Supplementary Figure 2. Unabridged biodistributions of Abraxane and albumin. Data for blood, lungs, liver, and spleen are reproduced from Figure 1 in the main text. (a) Comparison of Abraxane biodistributions at 1 mg/kg and 5 mg/kg bolus doses in naive mice and mice treated with intravenous bacterial lipopolysaccharides (LPS). (b) Biodistributions of albumin in naive mice and mice treated with intravenous LPS. (c) Biodistribution of 5 mg/kg Abraxane in mice afflicted with inhaled H1N1 virus. (d) Biodistributions of 5 mg/kg Abraxane in C3 knockout mice, either naive or treated with intravenous LPS.

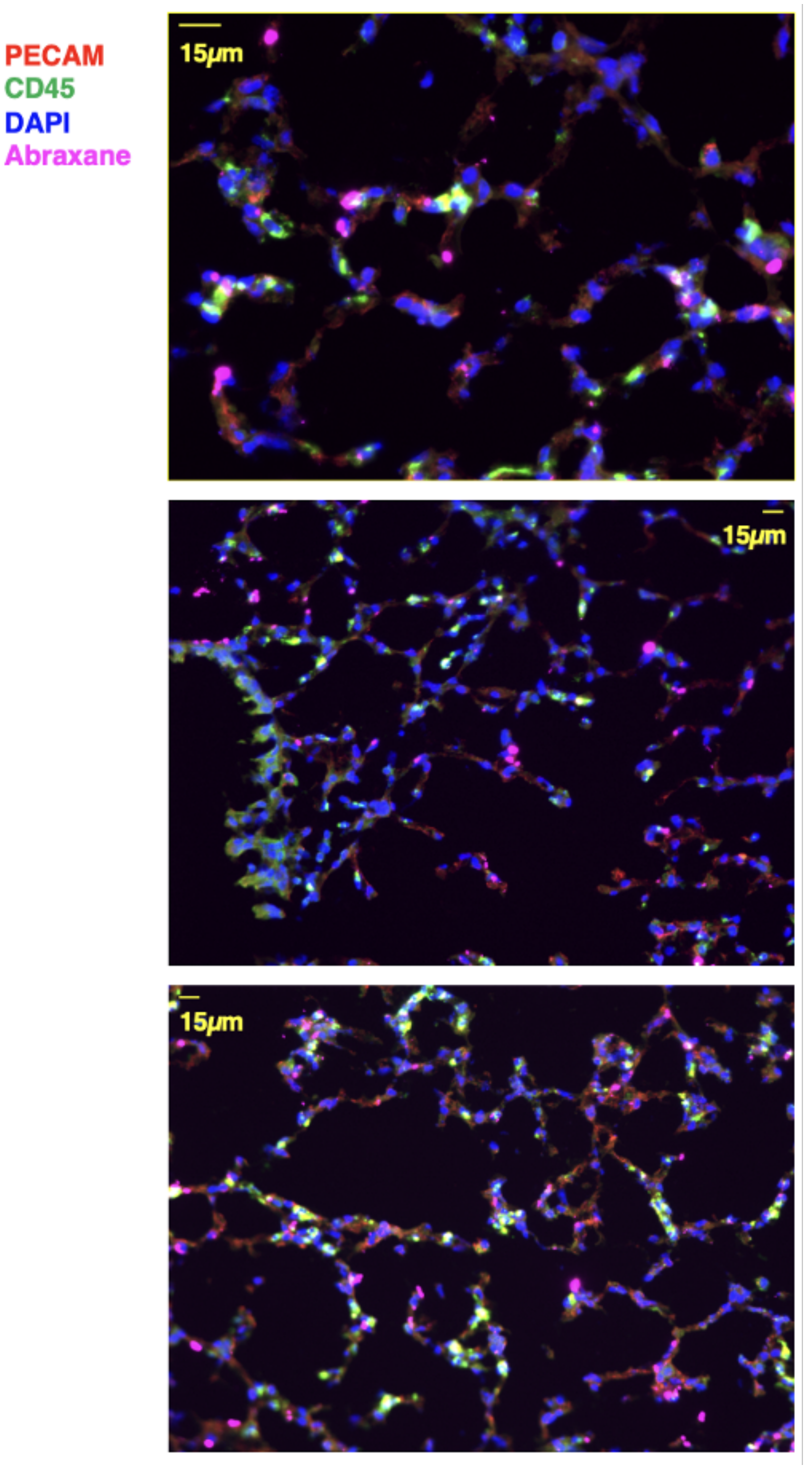


Supplementary Figure 3. Lung histology showing Abraxane uptake in intravenous LPS-treated mice. Data represent additional views of the staining depicted in Figure 2a in the main text, reflecting data from two mice. Bright punctate Abraxane signal, primarily in pulmonary vasculature, is evident across all images.

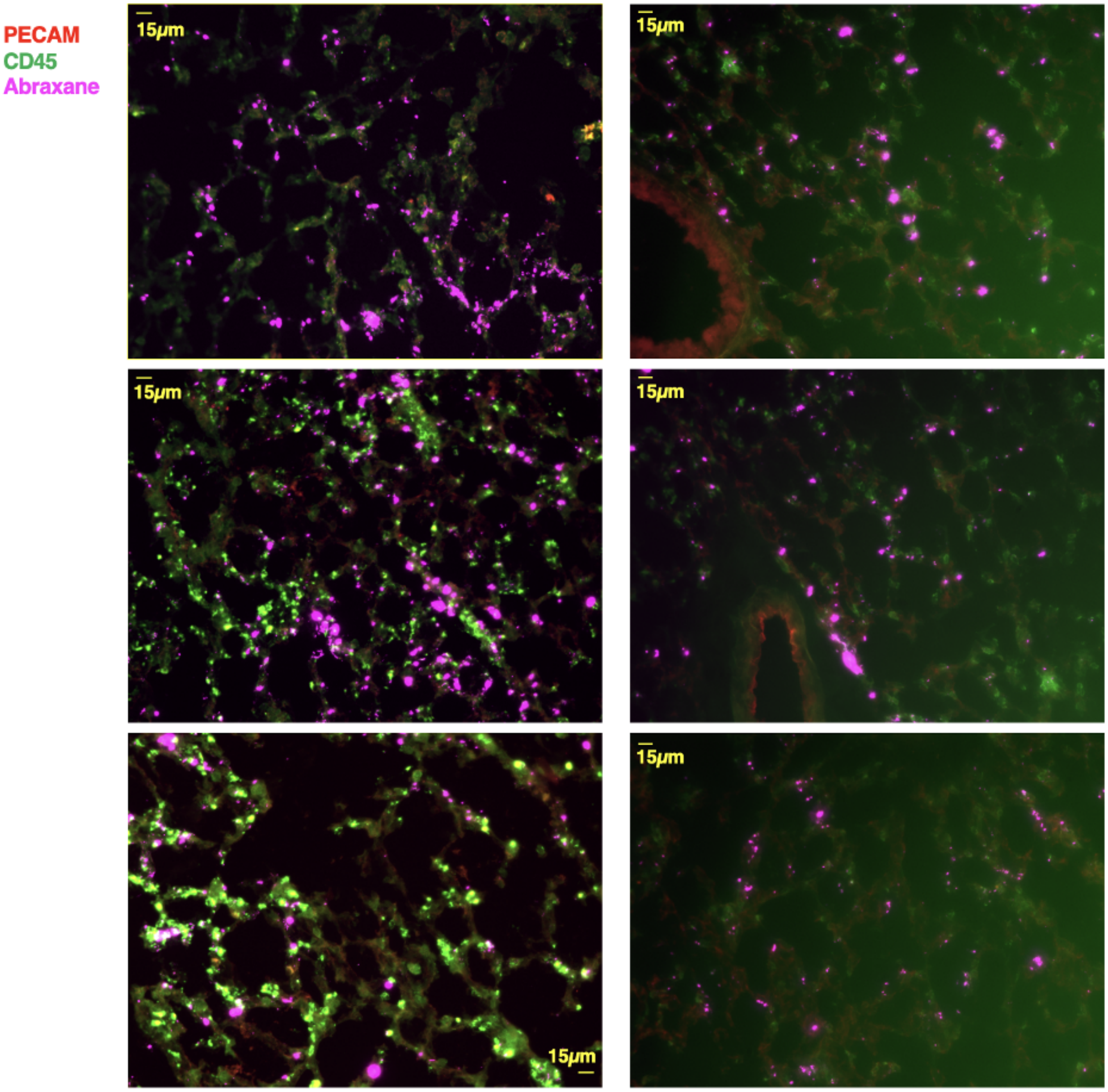


Supplementary Figure 4. Lung histology showing Abraxane uptake in intravenous LPS-treated mice. Data show micrographs from slides where DAPI staining was eliminated and CD45 staining was amplified with a secondary antibody to highlight Abraxane colocalization with leukocytes.

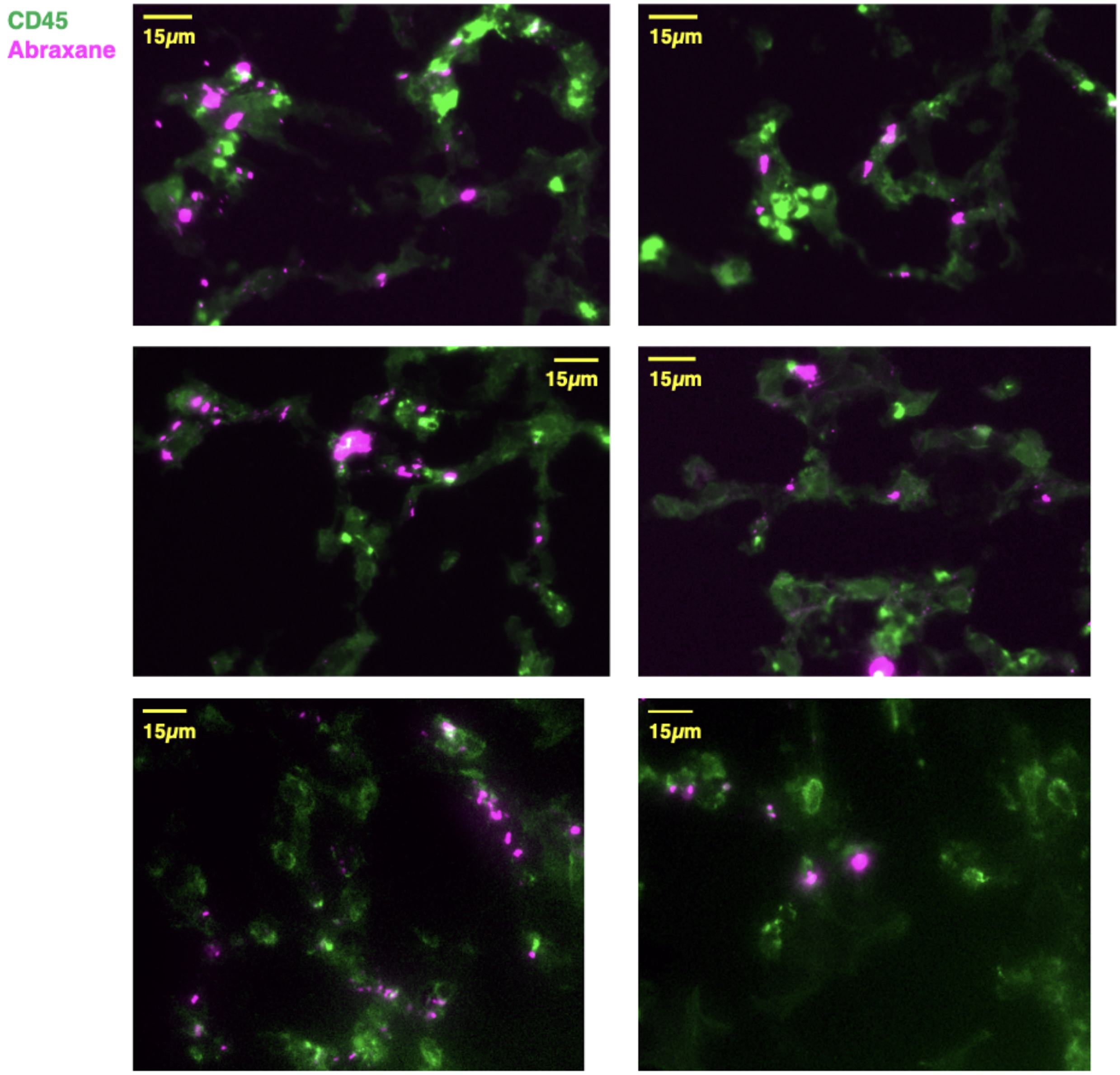


Supplementary Figure 5. Lung histology showing Abraxane uptake in leukocytes in intravenous LPS-treated mice. Data reflect magnified images from lung sections stained as in Supplementary Figure 4, with focal areas selected to show Abraxane accumulation in and around leukocytes.

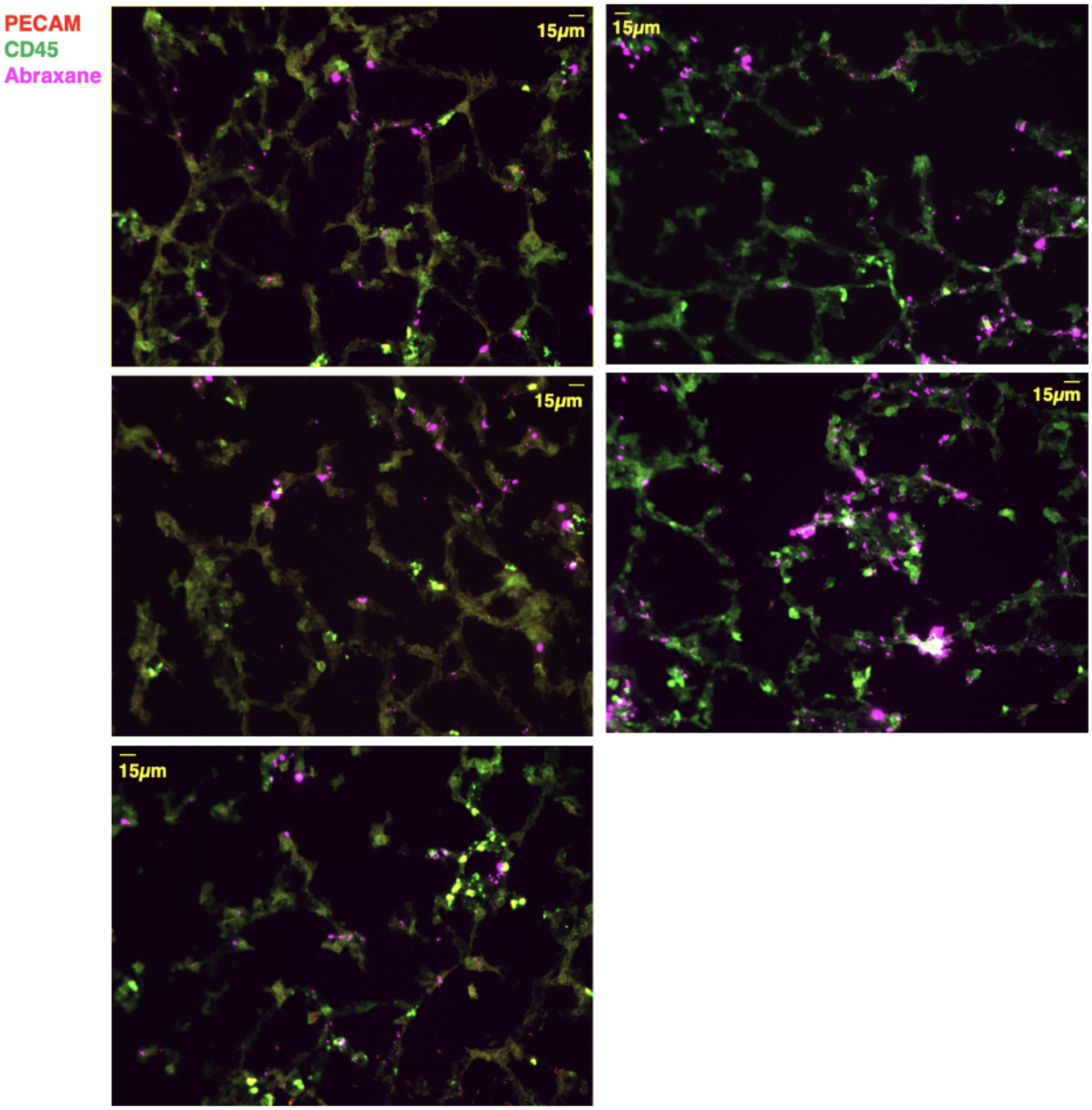


Supplementary Figure 6. Lung histology showing Abraxane uptake in naïve mice. Slides were stained as in Supplementary Figure 4 to emphasize leukocytes surrounding Abraxane signal.

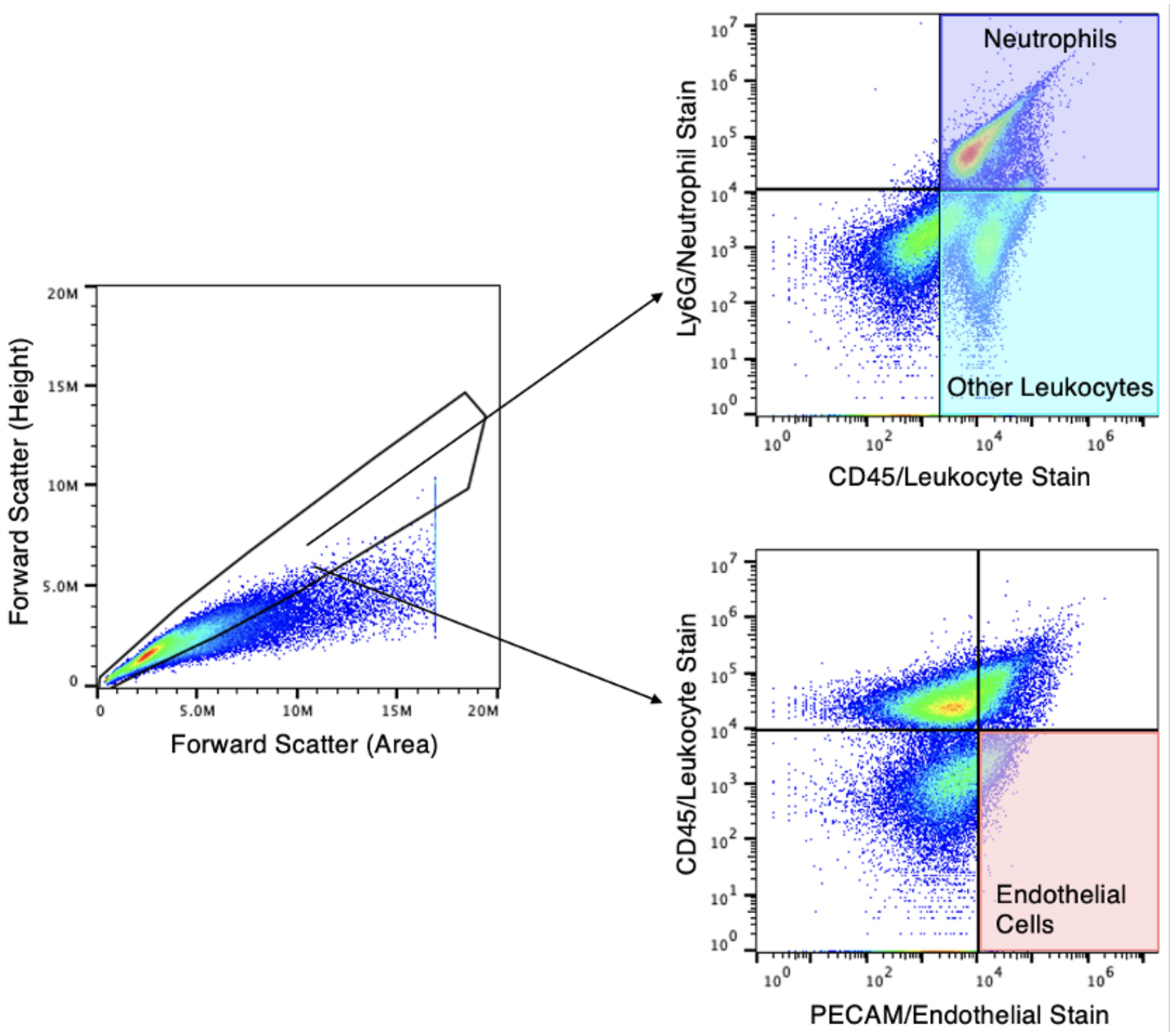


Supplementary Figure 7. Gating strategy for mouse lungs flow cytometry. Leftmost panel: All cells were analyzed for the presence of doublets by plotting forward scattering signal (height) vs. forward scattering signal (area), assessing doublets for inclusion or exclusion by events with disproportionate area for a given height. Upper right panel: Leukocytes were identified by an above-threshold CD45 staining and neutrophils were identified by an above-threshold Ly6G staining among CD45-positive events. Lower right panel: Among CD45-negative events, endothelial cells were identified by above-threshold PECAM staining. CD45, Ly6G, and PECAM staining thresholds were determined based on samples without staining, set such that unstained samples measured <1% stain-positive events.

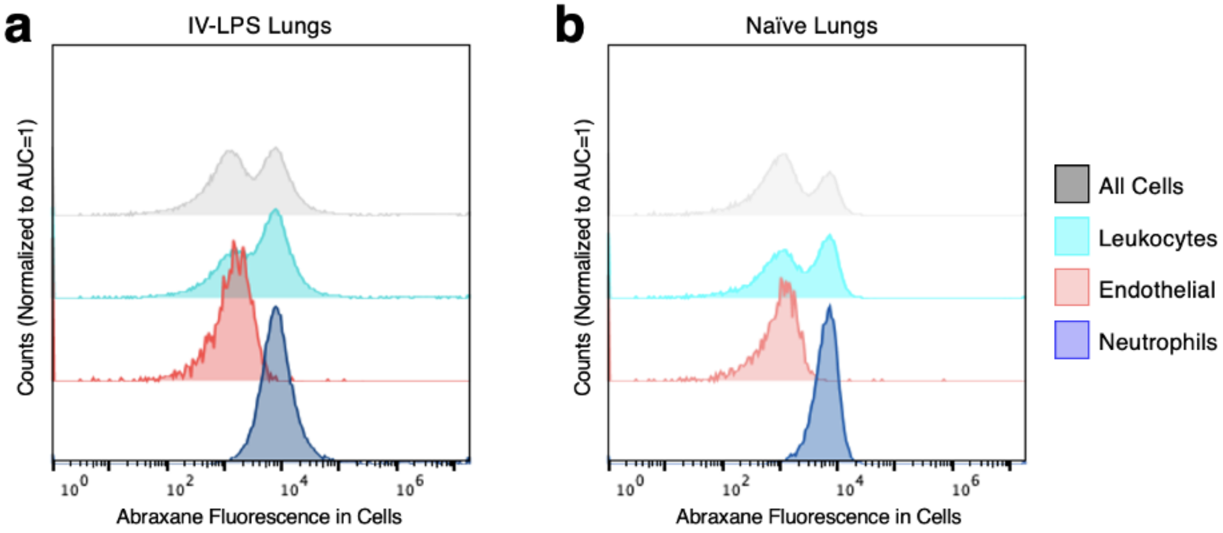


Supplementary Figure 8. Mouse lungs flow cytometry data showing Abraxane signal in different cell types (normalized to total abundance of each cell type) in lungs from mice treated with intravenous LPS (a) and naive lungs (b). Data are as in Figure 2c in the main text, but for different animals. Comparison of (a) and (b) shows the broad similarity of Abraxane targeting to neutrophils in naive vs. LPS-treated mice, even accounting for the different abundance and phenotypes of neutrophils under the two conditions.

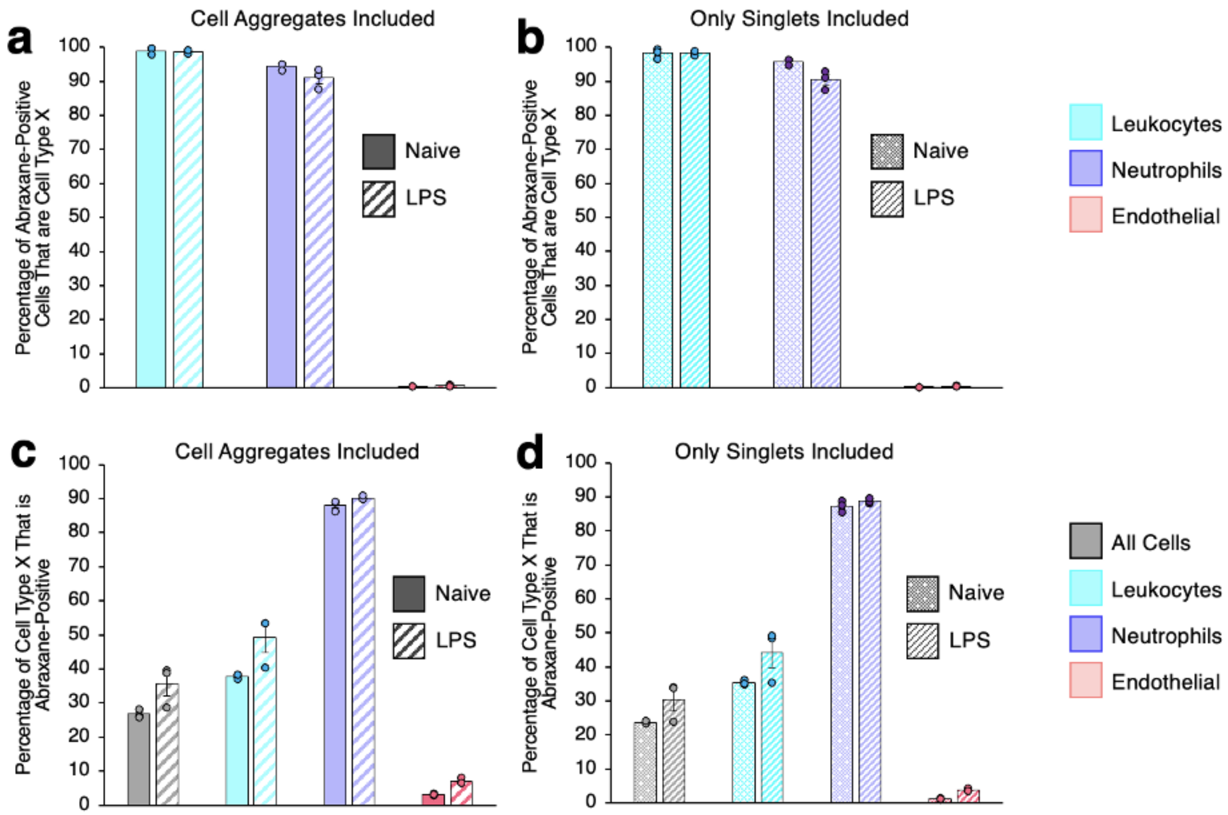


Supplementary Figure 9. Summary data showing Abraxane distribution in mouse lungs flow cytometry, including the role of cell aggregates vs. isolated cells. (a-b) Percentage of total Abraxane-positive cells in mouse lungs that are leukocytes, neutrophils, or endothelial cells. Data in (a) omit the singlet gating shown in Supplementary Figure 8 in order to introduce the effects of aggregated leukocytes to our data. Comparison of (a) and (b) shows no effects of aggregates in Abraxane uptake in neutrophils. (c-d) Percentage of all cells, all leukocytes, neutrophils, and endothelial cells that are Abraxane positive. As in (a), data in (c) omit singlet gating. Comparison of (c) and (d) shows no effects of cell aggregates in Abraxane selectivity for neutrophils.

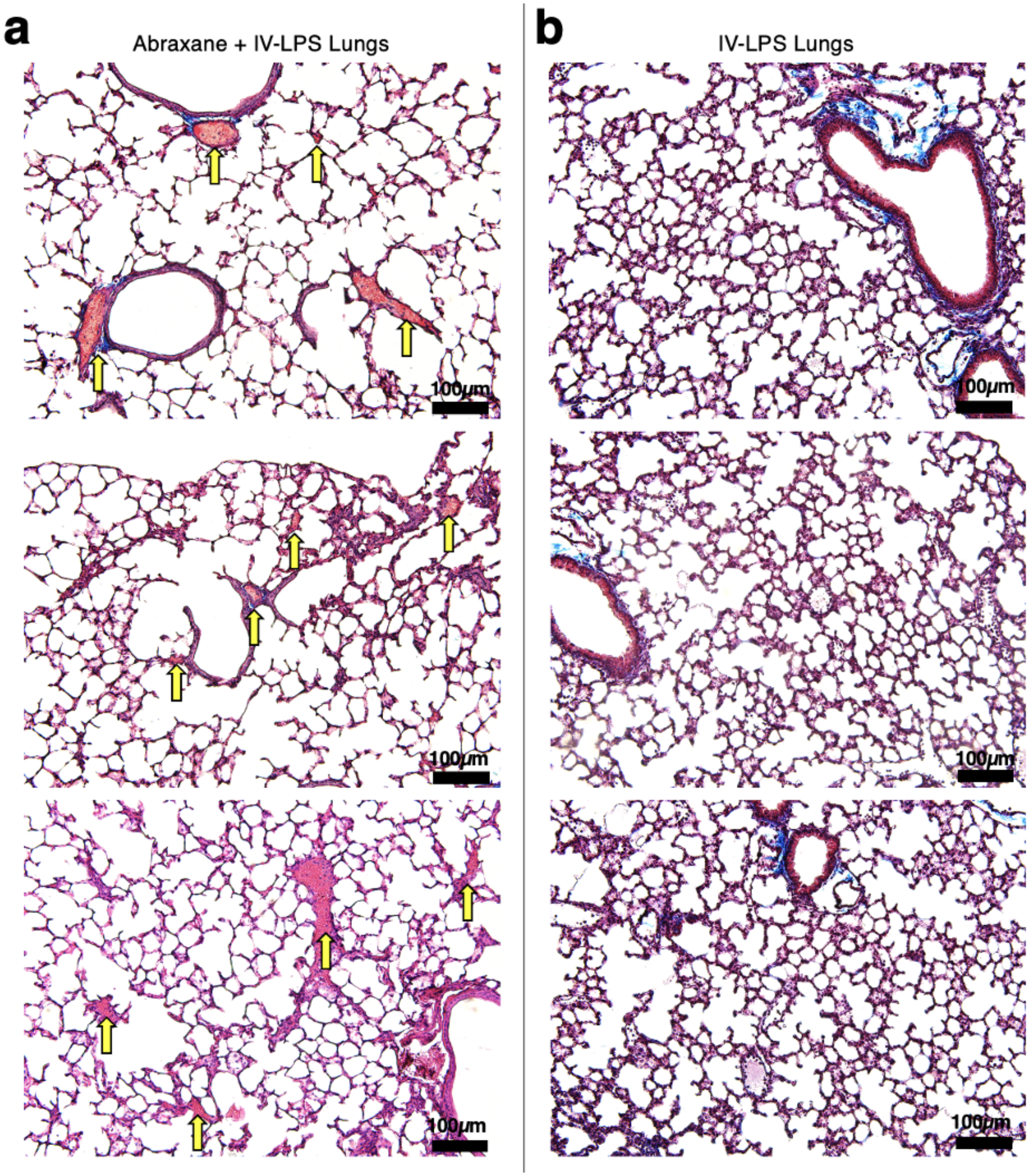


Supplementary Figure 10. Masson trichrome stained micrographs of lungs from mice treated with intravenous LPS, with (a) or without (b) Abraxane treatment. Yellow arrows point to occlusive thrombi in large vessels in (a).

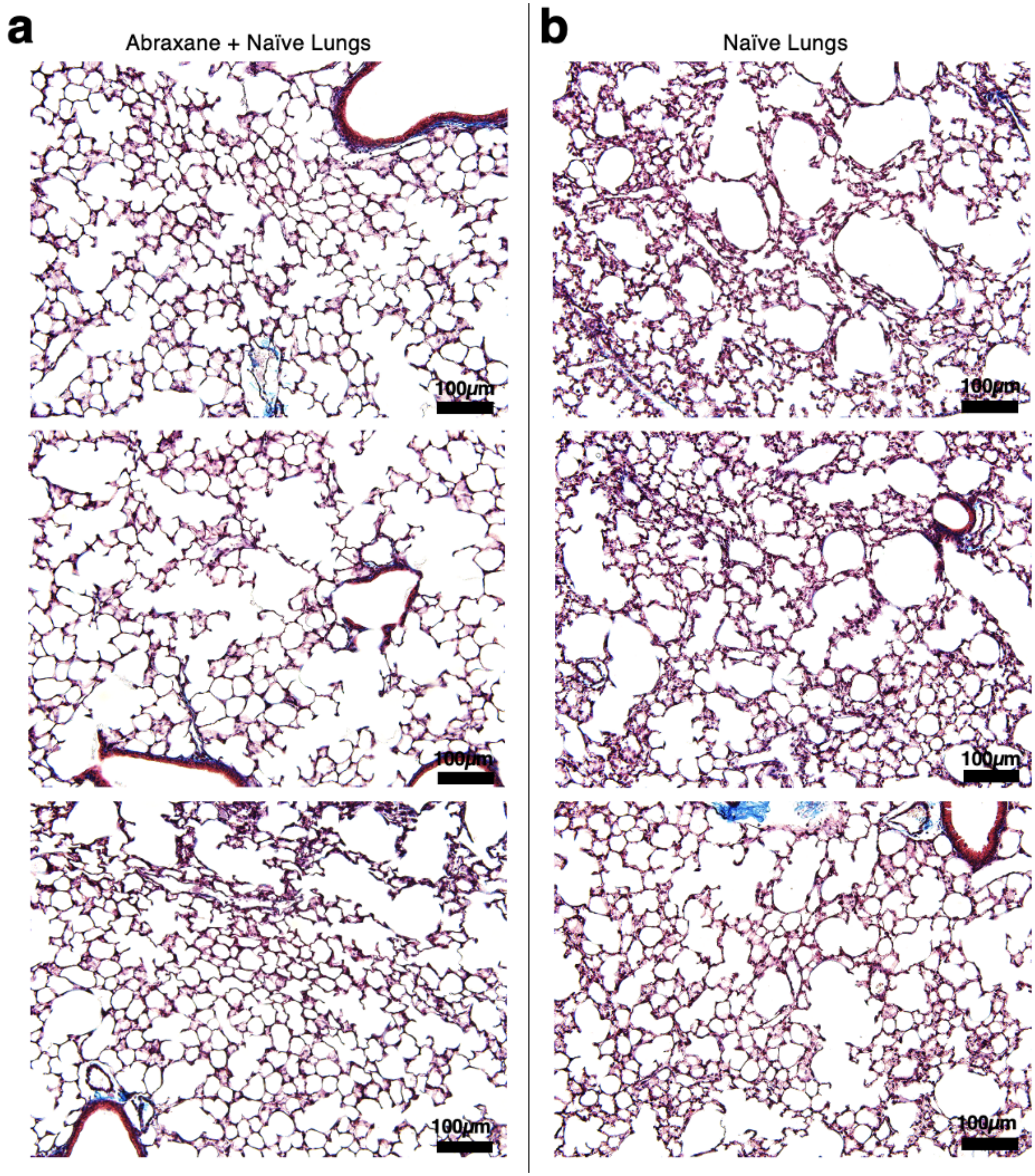


Supplementary Figure 11. Masson trichrome stained micrographs of naive mouse lungs, with (a) or without (b) Abraxane treatment.

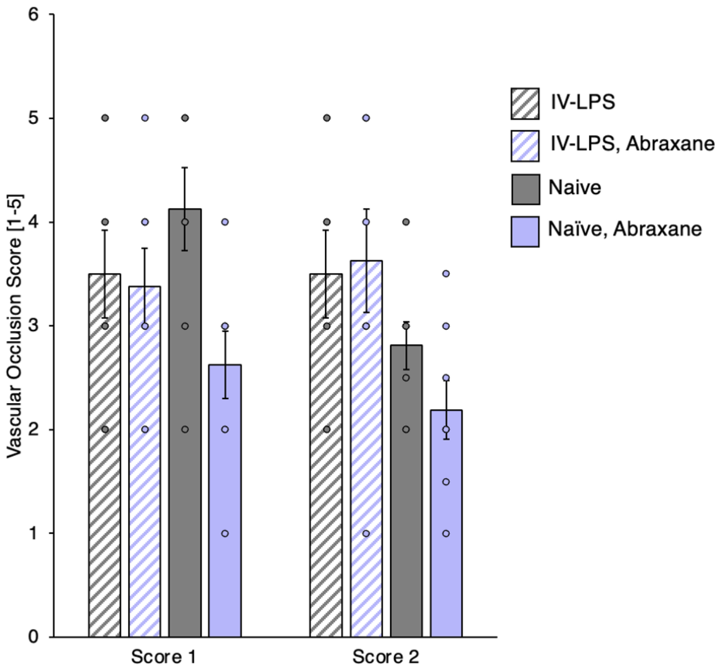


Supplementary Figure 12. Blinded scoring of Masson trichrome stained micrographs under conditions depicted in Figure 3a in the main text and Supplementary Figures 11 and 12, assessing total likelihood of vascular occlusion by thrombi or inflammatory infiltrates in all vessels.

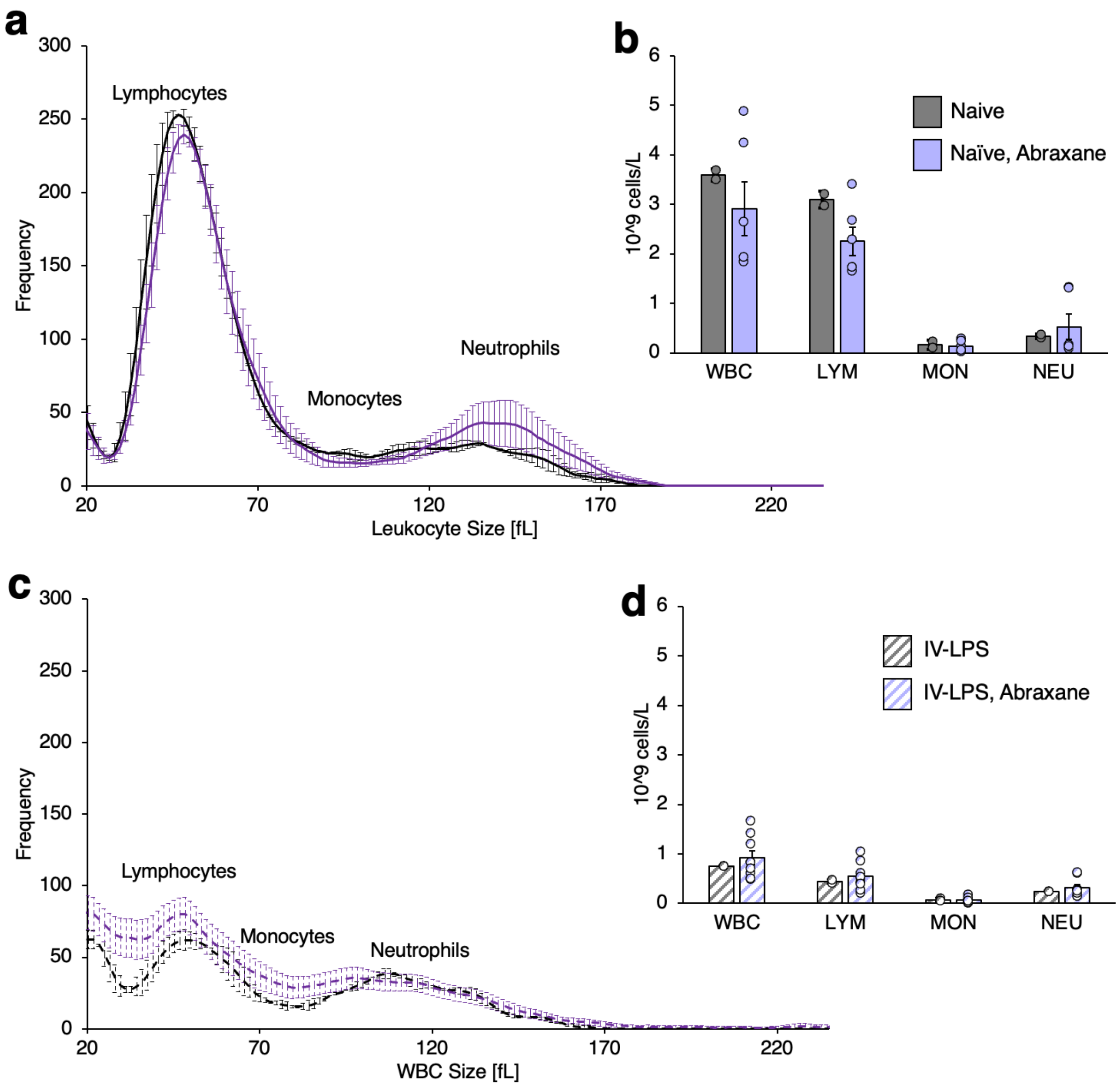


Supplementary Figure 13. Complete blood count data assessing Abraxane effects on circulating white blood cells. (a) Size distribution of circulating leukocytes in naïve mice and mice treated with 5 mg/kg Abraxane bolus 30 minutes prior to blood draw. (b) Counts of total leukocytes (WBC), lymphocytes (LYM), monocytes (MON), and neutrophils (NEU) derived from data in (a). (c) Size distribution of circulating leukocytes in mice treated with intravenous LPS (5.5 hours prior to blood draw) and mice treated with intravenous LPS followed 5 hours later by 5 mg/kg Abraxane bolus. (d) Counts of total leukocytes, lymphocytes, monocytes, and neutrophils derived from data in (c).

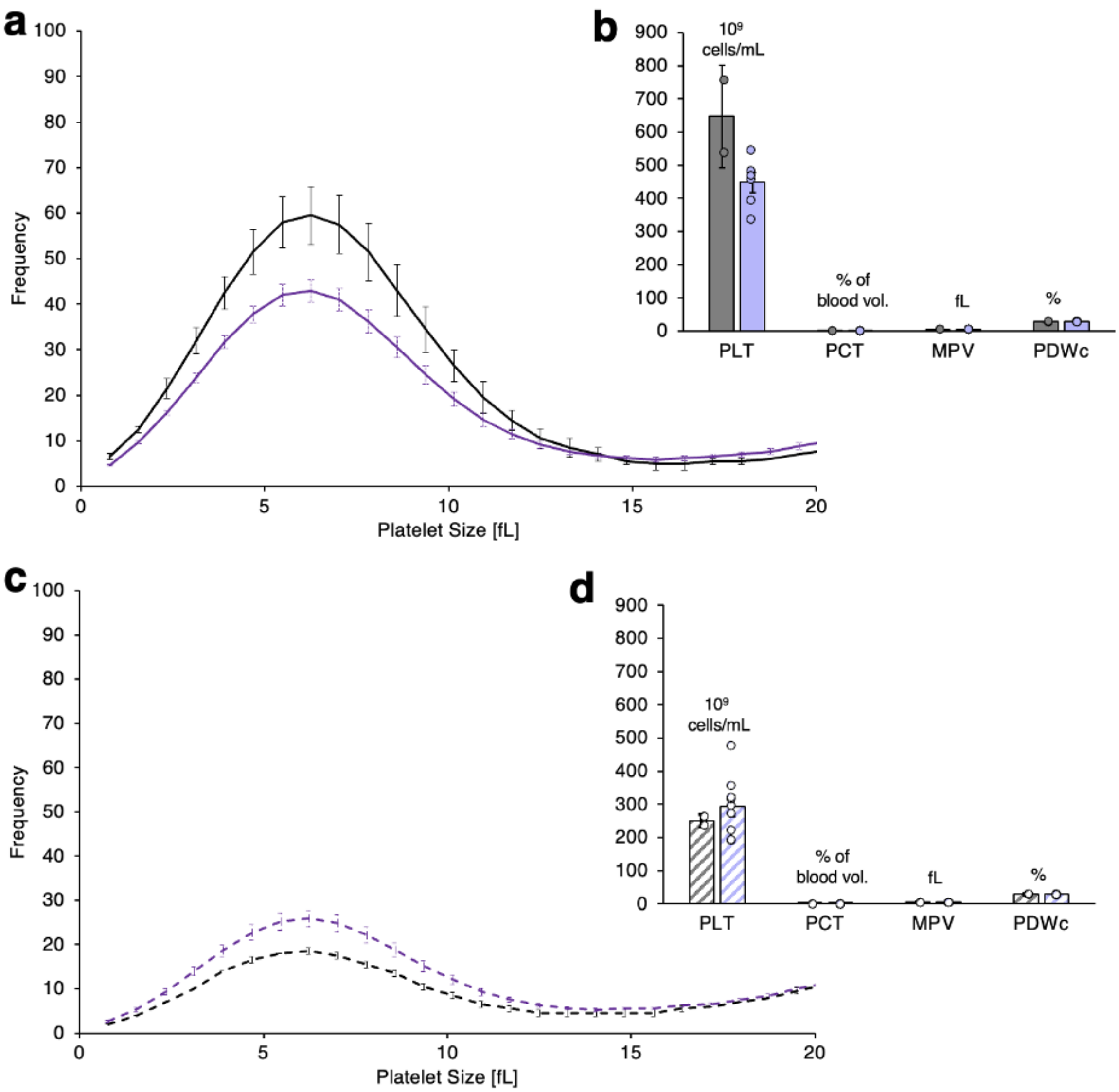


Supplementary Figure 14. Complete blood count data assessing Abraxane effects on circulating platelets. (a) Size distribution of circulating platelets in naïve mice and mice treated with 5 mg/kg Abraxane bolus 30 minutes prior to blood draw. (b) Platelet count (PLT), platelet volume fraction in blood (PCT), mean platelet volume (MPV), and deviation in platelet volume (PDWc) derived from data in (a). (c) Size distribution of circulating platelets in mice treated with intravenous LPS (5.5 hours prior to blood draw) and mice treated with intravenous LPS followed 5 hours later by 5 mg/kg Abraxane bolus. (d) Platelet count, platelet volume fraction in blood, mean platelet volume, and deviation in platelet volume derived from data in (c).

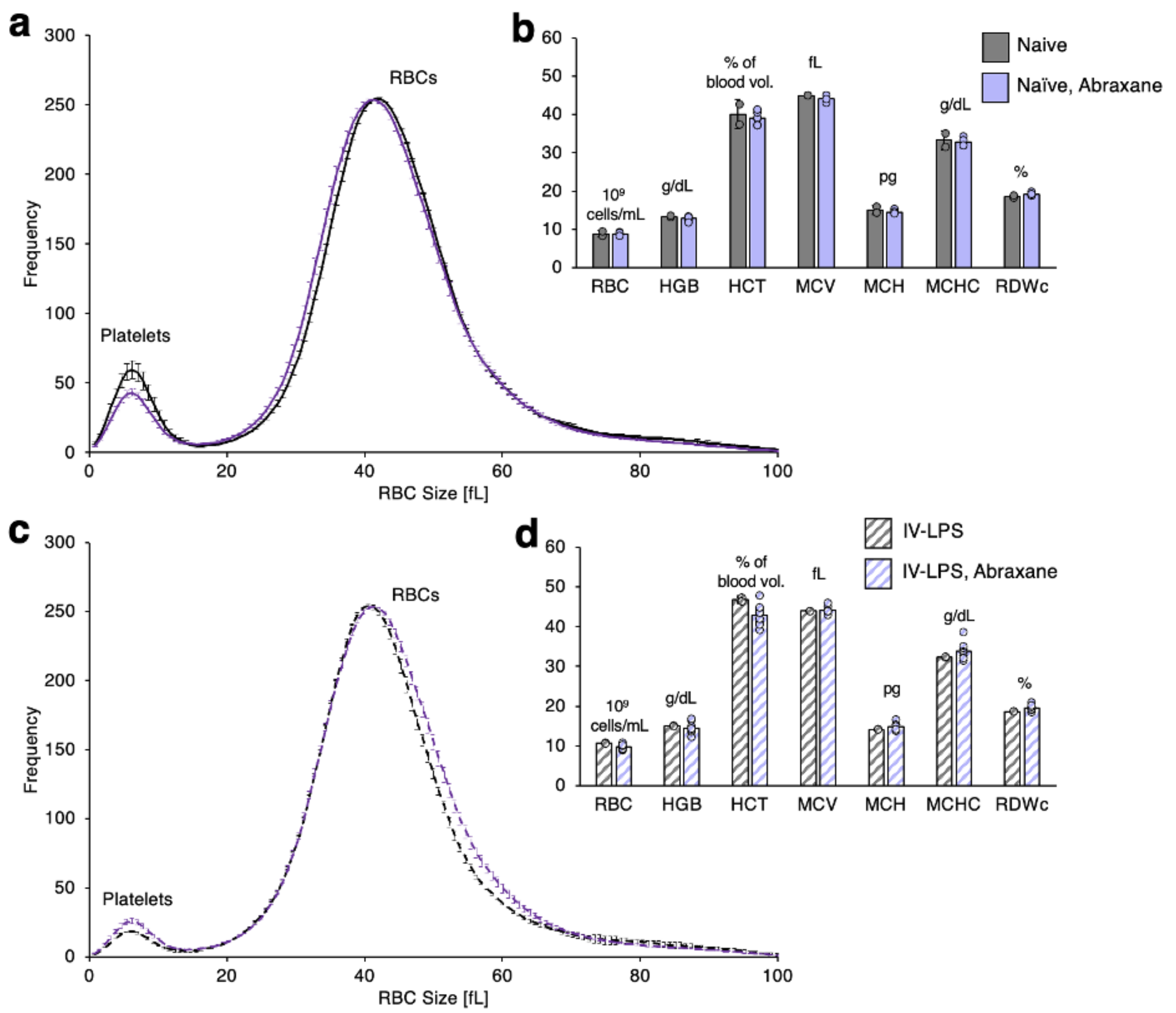


Supplementary Figure 15. Complete blood count data assessing Abraxane effects on circulating red blood cells. (a) Size distribution of circulating red blood cells in naïve mice and mice treated with 5 mg/kg Abraxane bolus 30 minutes prior to blood draw. (b) Red blood cell count (RBC), hemoglobin concentration in blood (HGB), red blood cell volume fraction in blood (HCT), mean red blood cell volume (MCV), mean red blood cell hemoglobin content (MCH), mean red blood cell hemoglobin concentration (MCHC), and deviation in red blood cell volume (RDWc) derived from data in (a). (c) Size distribution of circulating red blood cells in mice treated with intravenous LPS (5.5 hours prior to blood draw) and mice treated with intravenous LPS followed 5 hours later by 5 mg/kg Abraxane bolus. (d) Red blood cell count, hemoglobin concentration in blood, red blood cell volume fraction in blood, mean red blood cell volume, mean red blood cell hemoglobin content, mean red blood cell hemoglobin concentration, and deviation in red blood cell volume derived from data in (c).

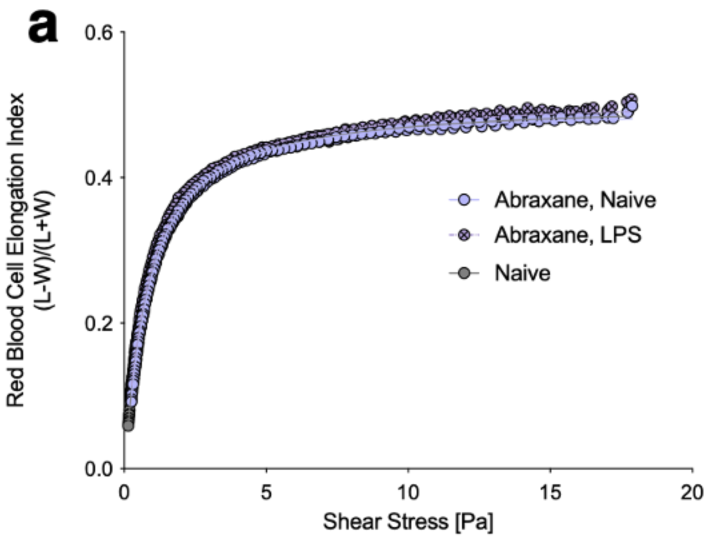


Supplementary Figure 16. Ektacytometry data assessing Abraxane effects on the mechanical properties of circulating red blood cells. (a) Stress-strain curves showing relative elongation of red blood cells in microfluidic devices as a function of applied shear stress. (b) Relative red blood cell elongation at 5 Pa and 10 Pa applied shear stress, alongside extrapolated maximum elongation at infinite shear.

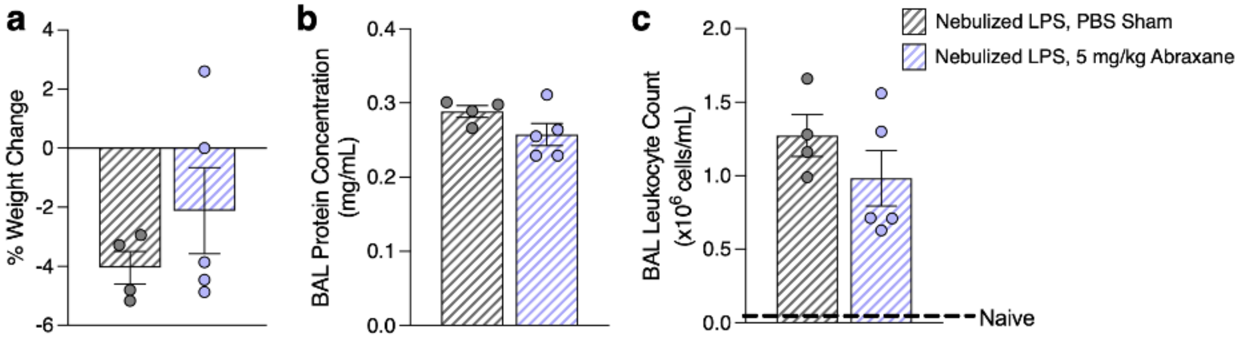


Supplementary Figure 17. Abraxane effects on the outcome of mouse model acute respiratory distress syndrome. (a) Change in body weight following treatment with nebulized LPS and either PBS sham or 5 mg/kg Abraxane bolus. (b) Protein concentration in bronchoalveolar lavage (BAL) fluid, as a measure of lung edema for conditions as in (a). (c) Leukocyte counts in BAL fluid, as a measure of inflammatory response in the lungs for conditions as in (a).

| Lung ID | Age | Sex | Cause of Death | Lobe Obtained/Perfused | Edema (Visual Inspection) | Hepatization | Smoking Damage |
|---|---|---|---|---|---|---|---|
| Radiotracer 1 | 29 | F | Anoxia | Right Upper | No | No | No |
| Radiotracer 2 | 28 | M | Anoxia, Cardiovascular | Left Upper | No | Left Lower | No |
| Radiotracer 3 | 45 | F | Cerebrovascular Accident | Left Upper | No | Left Lower | Mild |
| Radiotracer 4 | 31 | M | Anoxia, Cardiovascular | Left Upper | No | Left Lower | Mild |
| Flow Cytometry/Histology 1 | 43 | F | Cerebrovascular Accident | Right Lower | No | Right Mid | Mild-Moderate |
| Flow Cytometry/Histology 2 | 42 | M | Cerebrovascular Accident | Left Lower | No | Left Lower | No |

Supplementary Figure 18. De-identified patient data describing rejected donor lungs used in radiotracing and flow cytometry/histology studies.

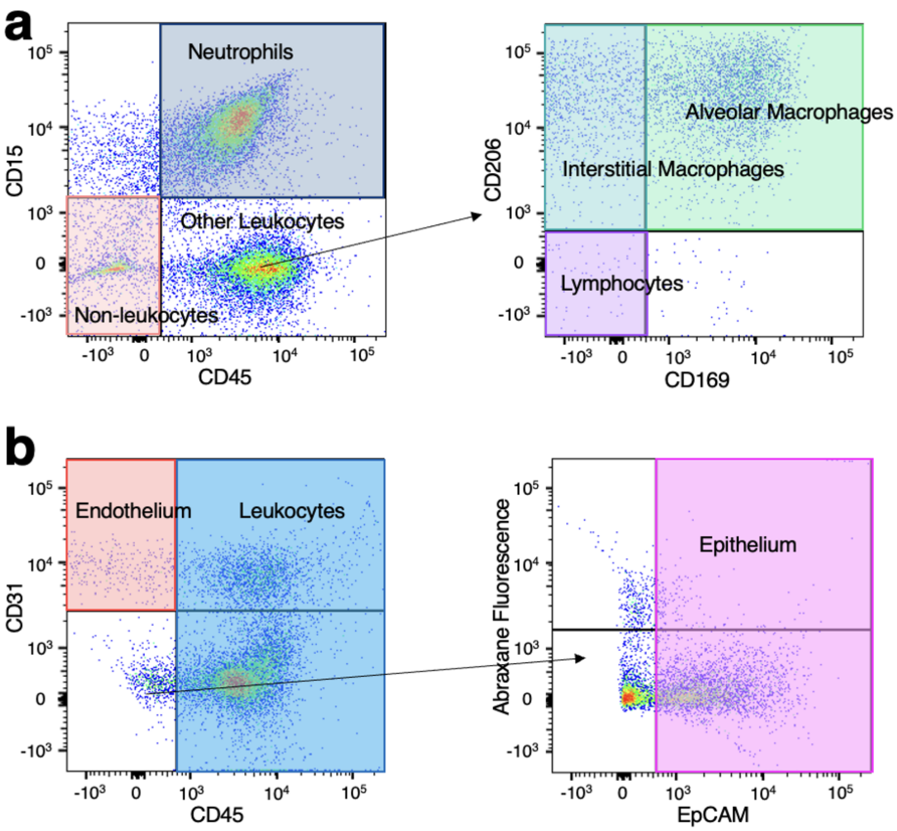


Supplementary Figure 19. Gating strategy for human lungs flow cytometry. (a) Gating strategy for staining panel used to identify leukocyte subtypes. (b) Gating strategy for staining panel used to identify endothelial cells and epithelial cells.

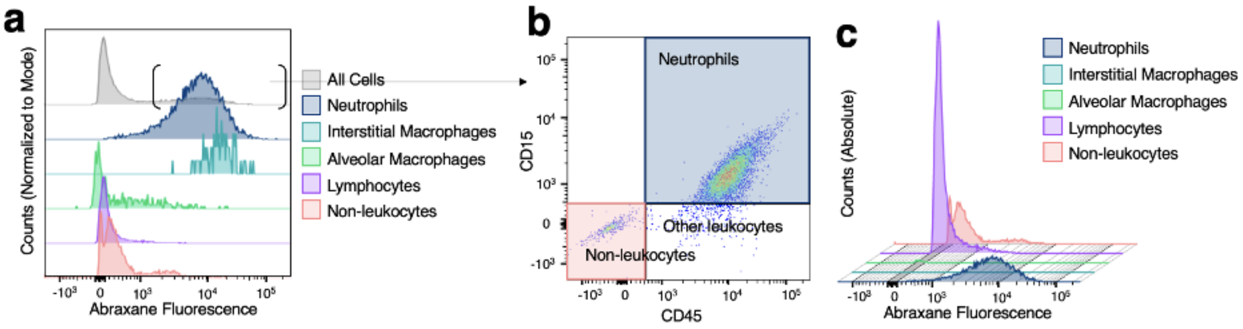


Supplementary Figure 20. Counterpart data for Figure 4c-e in the main text, showing findings for a second lung donor. (a) Normalized histograms of Abraxane fluorescence in different cell types. (b) Among all Abraxane positive cells, staining for CD15 and CD45 to identify Abraxane-positive neutrophils and other leukocytes. (c) Non-normalized histograms showing true frequency of Abraxane fluorescence in different cell types.

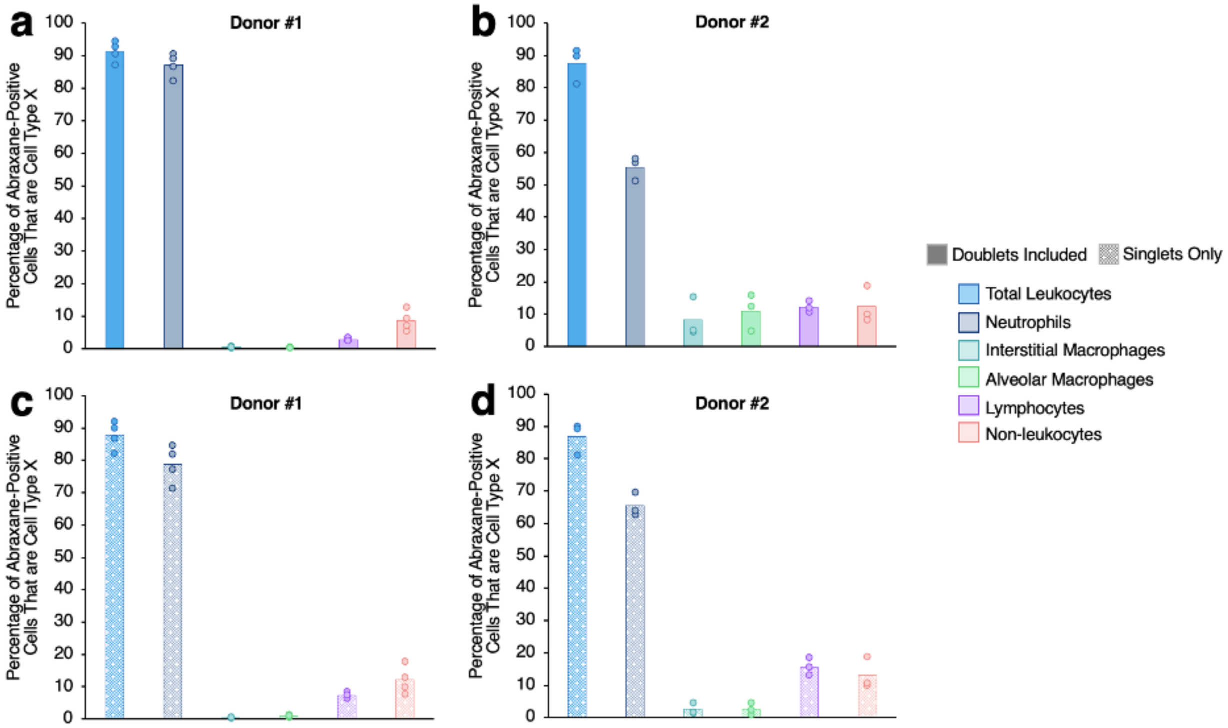


Supplementary Figure 21. Summary data showing Abraxane distribution to different cell types in human lungs flow cytometry, including the role of cell aggregates vs. isolated cells.

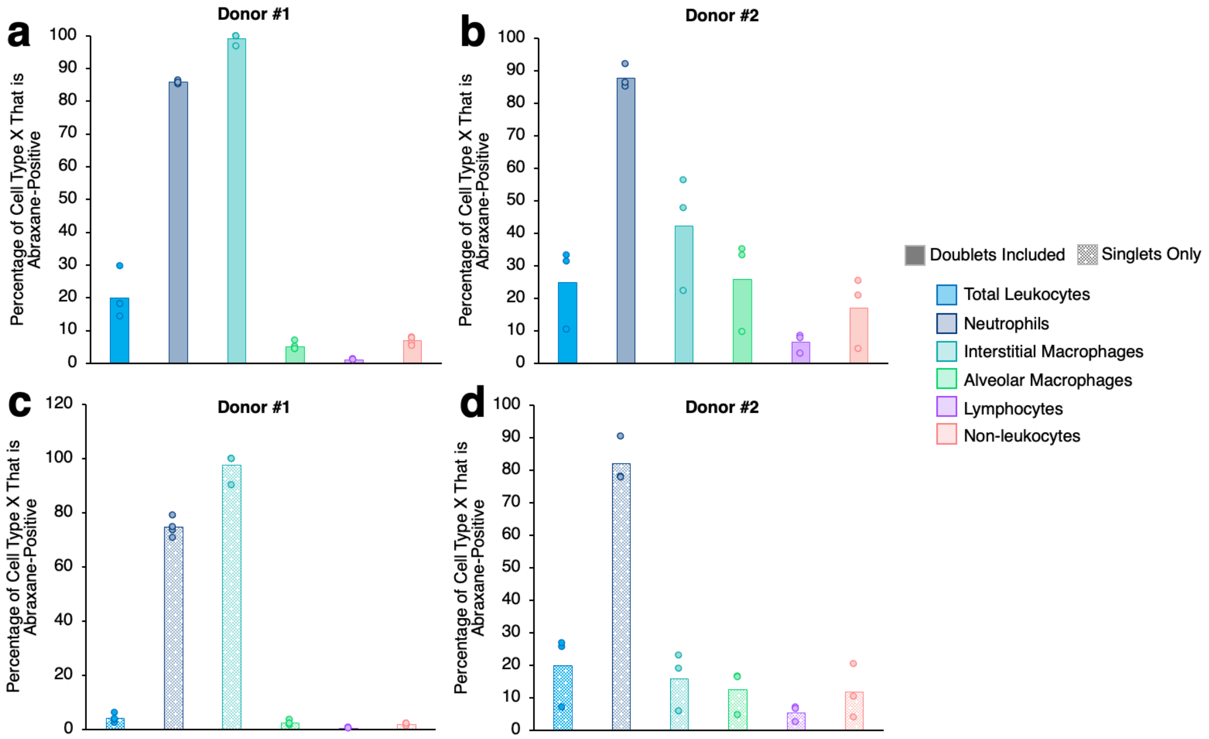


Supplementary Figure 22. Summary data showing Abraxane uptake in different cell types in human lungs flow cytometry, including the role of cell aggregates vs. isolated cells.

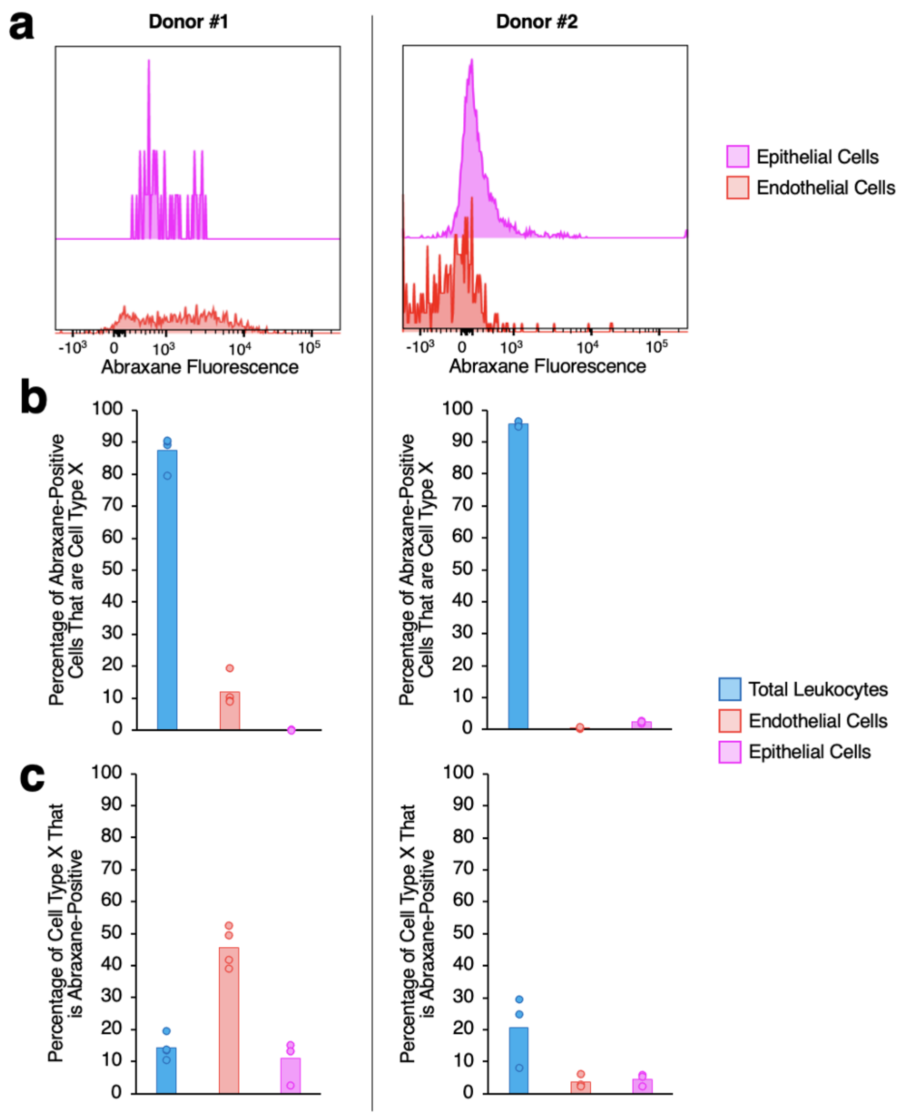


Supplementary Figure 23. Human lungs flow cytometry data tracking Abraxane uptake in endothelial cells and epithelial cells.

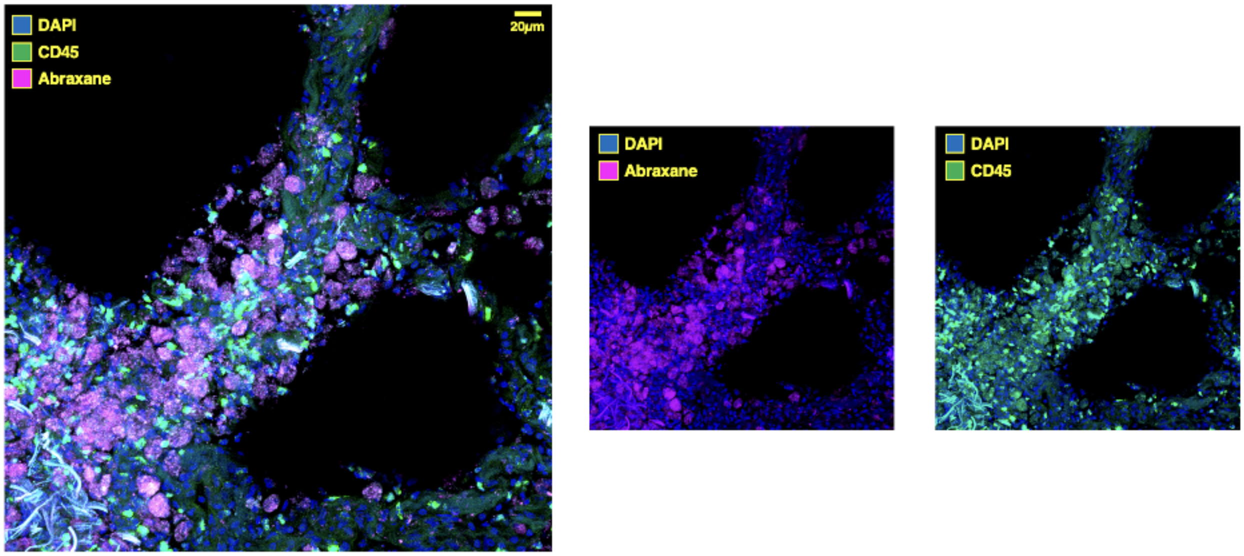


Supplementary Figure 24. Histology of human lungs showing Abraxane colocalization with leukocytes.

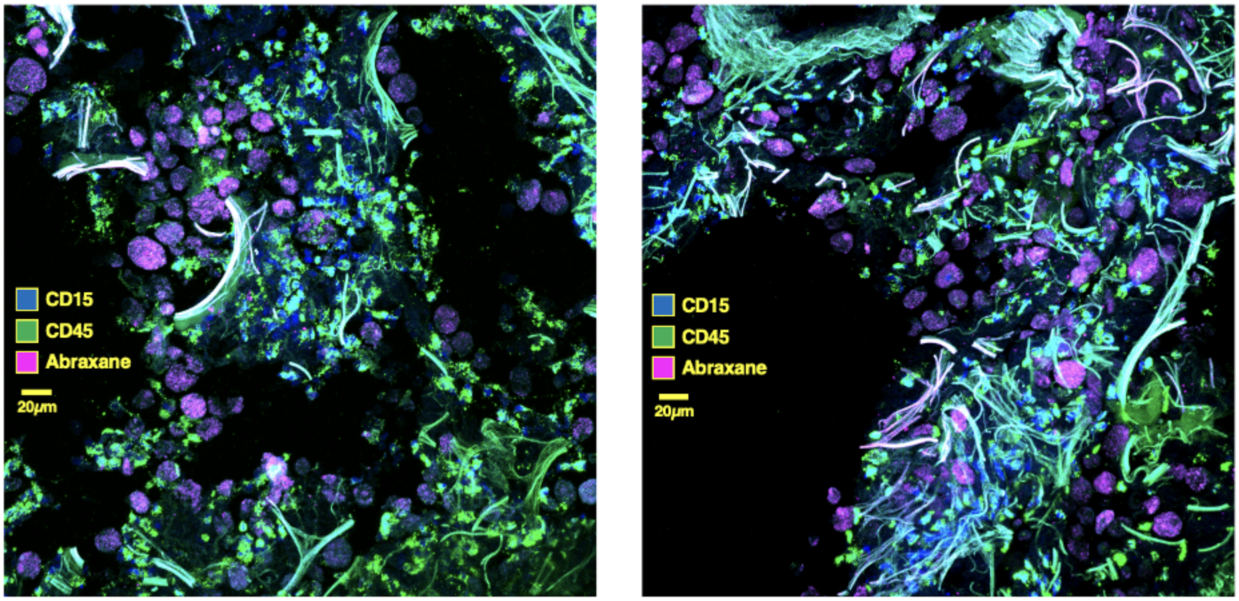


Supplementary Figure 25. Histology of human lungs showing Abraxane colocalization with neutrophils.

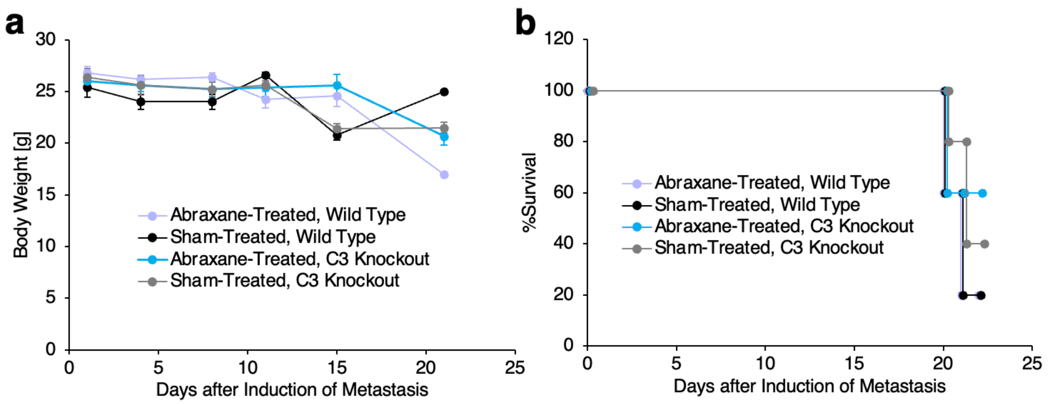


Supplementary Figure 26. Effects on body weight (a) and survival (b) for Abraxane treatment in wild type and C3 knockout mice with lung metastases.

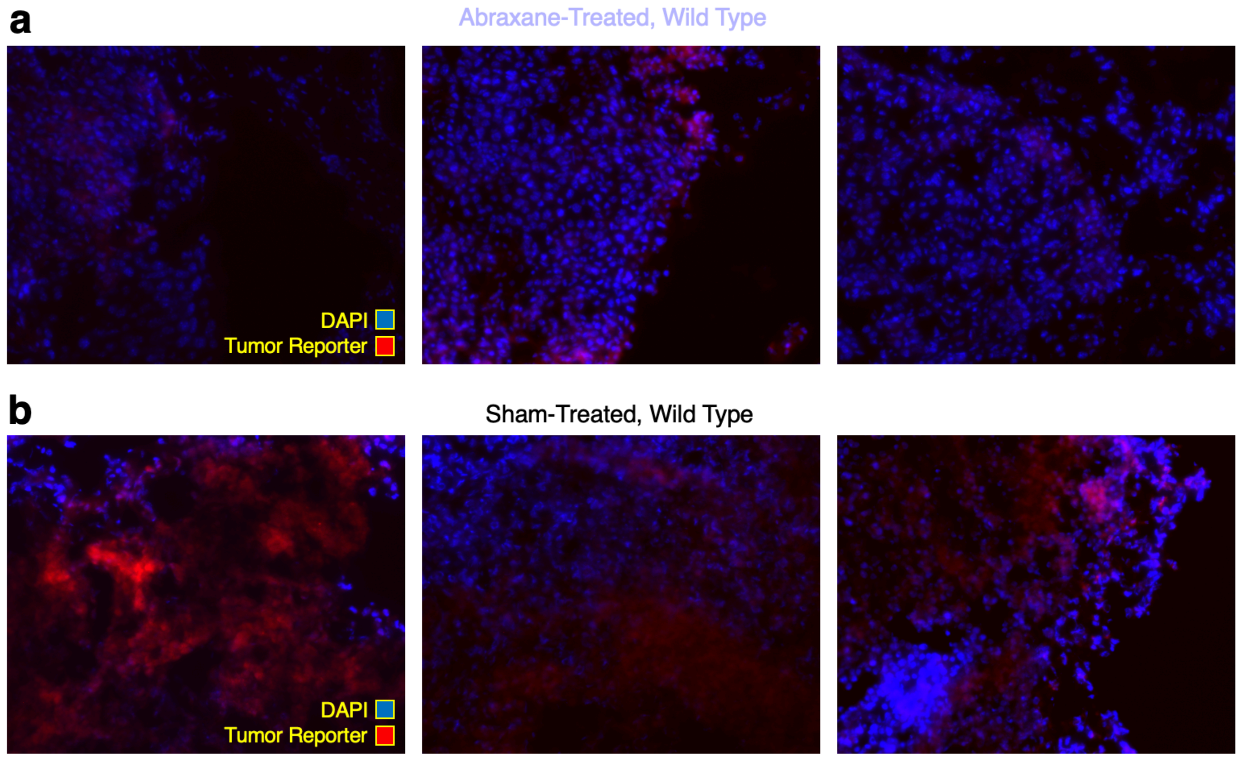


Supplementary Figure 27. Histology indicating tumor reporter fluorescence as a function of Abraxane treatment in wild type mice. (a) Representative images from three mice treated with Abraxane five times over ten days following induction of lung metastases, followed by lung harvesting at 21 days. (b) Representative images as in (a) but for mice treated with saline sham rather than Abraxane.

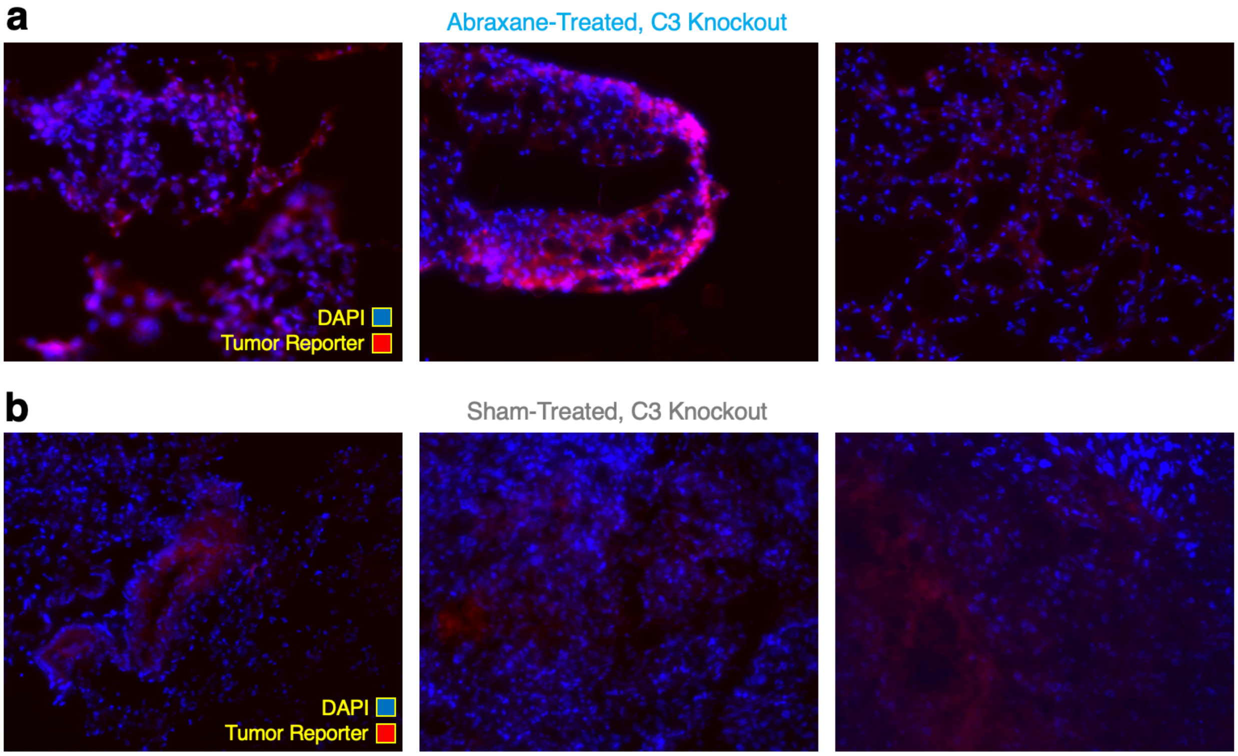


Supplementary Figure 28. Histology indicating tumor reporter fluorescence as a function of Abraxane treatment in C3 knockout mice. (a) Representative images from three C3 knockout mice treated with Abraxane five times over ten days following induction of lung metastases, followed by lung harvesting at 21 days. (b) Representative images as in (a) but for C3 knockout mice treated with saline sham rather than Abraxane.

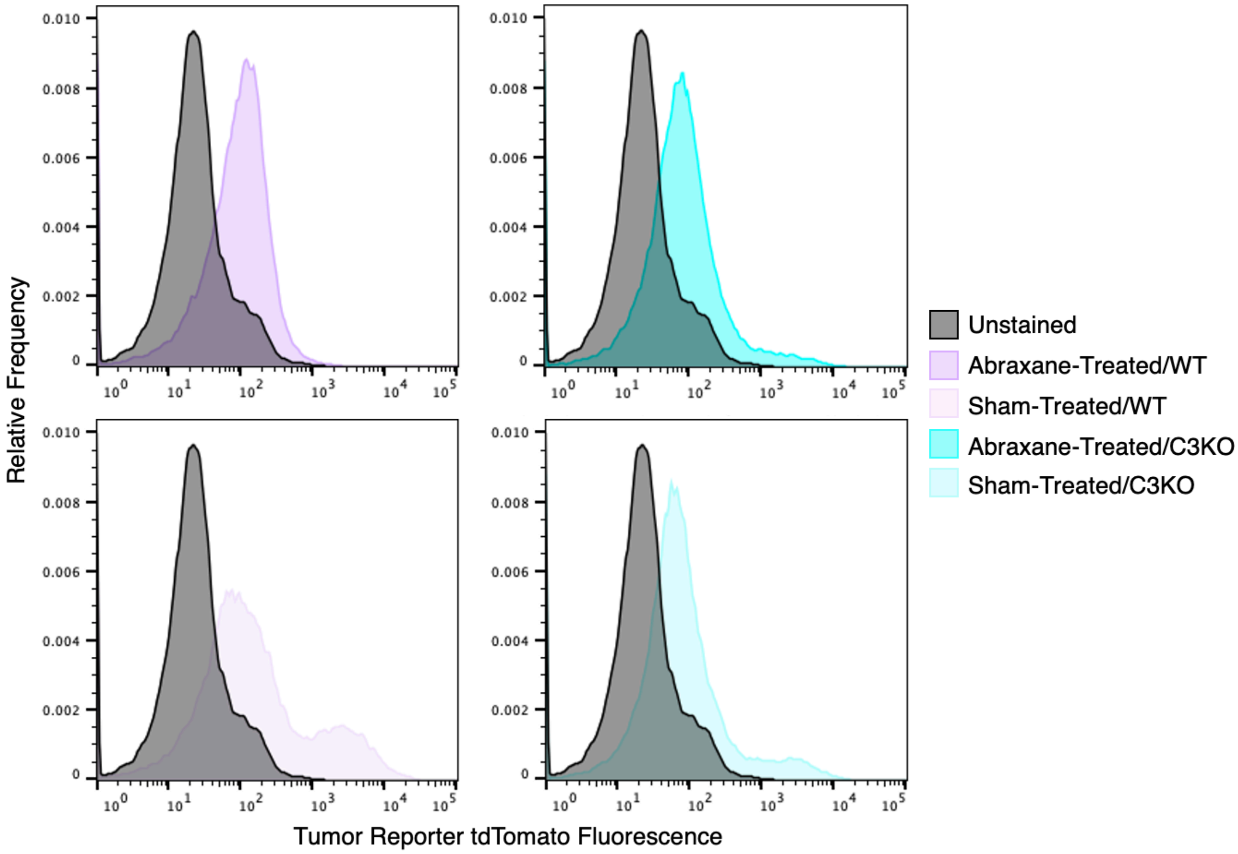


Supplementary Figure 29. Flow cytometry data indicating tumor reporter fluorescence as a function of Abraxane treatment in single cell suspensions prepared from lungs of wild type and C3 knockout mice with lung metastases. Representative histograms depicting relative frequency of cells vs. tumor reporter tdTomato fluorescence intensity in the cells. Data is normalized such that area under curve = 1 for each histogram. Leftmost panels show data for single cell suspensions prepared from wild type lungs, with top = Abraxane-treated, lower = sham-treated. Rightmost panels show data for C3 knockout lungs, with top = Abraxane-treated, lower = sham-treated. A histogram for a single cell suspension from a mouse receiving no B16-F10 reporter cells is shown in gray in each panel. tdTomato reporter-positive cells were assessed via a gate excluding 98% of cells from the unstained control mouse histogram.

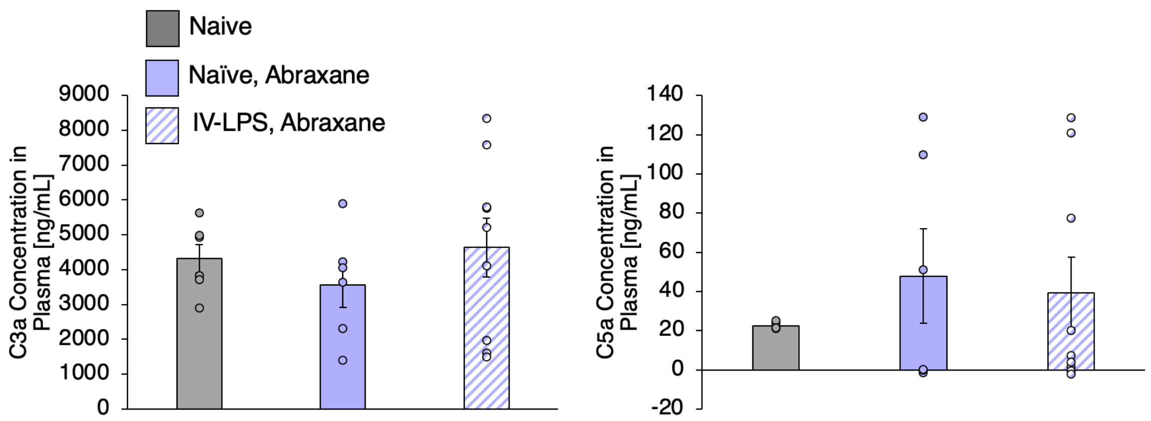


Supplementary Figure 30. Levels of complement fragments C3a and C5a in mouse plasma 30 minutes after treatment with 5 mg/kg bolus Abraxane. For mice receiving intravenous LPS prior to Abraxane dosing, Abraxane was administered 5 hours after LPS.
